# Branching Varies with Light Limitation Scenarios in relation with Changes in Carbon Source-Sink Dynamics

**DOI:** 10.64898/2026.01.27.702021

**Authors:** Anne Schneider, Frédéric Boudon, Sabine Demotes-Mainard, Lydie Ledroit, Maria-Dolores Perez-Garcia, Cédric Cassan, Yves Gibon, Christophe Godin, Soulaiman Sakr, Jessica Bertheloot

**Affiliations:** Univ Angers, Institut Agro, INRAE, IRHS, SFR QUASAV, F-49000 Angers, France; CIRAD, UMR AGAP Institut, 34398 Montpellier, France; AGAP Institut, CIRAD, INRAE, Institut Agro, Université de Montpellier, Montpellier, France; INRAE, Univ. Bordeaux, UMR 1332, Biologie du Fruit et Pathologie, 33882 Villenave d’Ornon, France; Laboratoire Reproduction et Developpement des Plantes, University of Lyon, ENS de Lyon, UCB Lyon 1, CNRS, INRA, Inria, F-69342, Lyon, France; LEPSE, Univ Montpellier, INRAE, Institut Agro Montpellier, Montpellier, France

**Keywords:** Architecture, Branching, Bud outgrowth, Light, Modelling, Plasticity, Source-Sink, Sugar

## Abstract

Bud outgrowth is a major component of plant architectural plasticity and is influenced by light conditions. While the inhibitory effect of low light intensity on branching is well documented, the underlying regulators remain debated and, especially, the role of sugar availability has never been thoroughly evaluated. Here, we combined experiments with a computational approach quantifying carbon source-sink balance in single-axis rose plants to investigate how continuous and transient light limitation regulate bud outgrowth. Continuous low light reduced photosynthesis, leading to decreased sugar availability and inhibited bud outgrowth. In contrast, a transient period of low light followed by high light unexpectedly stimulated bud outgrowth, shortened the delay between outgrowth of successive buds, and produced an over-branched phenotype. This response resulted from a non-reversible reduction in the growth of apical organs appearing under low light, which lowered carbon demand and caused sugar over-accumulation after the return to high light. Manipulating carbon supply and demand through leaf masking, photosynthetic inhibition, and targeted sucrose feeding causally confirmed the central role of sugar availability in these contrasting responses. Beyond these findings, key requirements for models simulating branching plasticity were identified and this work provides a basis for predicting branching responses under fluctuating and complex light environments.

**Highlight:** Bud outgrowth, a key component of plant plasticity, is regulated by light intensity through sugar availability. Continuous and transient low light have opposite effects by limiting sugar production and use, respectively.

## Introduction

As sessile organisms, plants exhibit high developmental plasticity, enabling them to adapt to their environment. Axillary bud outgrowth is a major component of this plasticity: buds are initiated at each leaf axil and may either grow immediately or enter dormancy that is released when conditions become favorable, thereby generating new axes and enabling space colonization (Barthelemy and Caraglio, 2007; Lang *et al*., 1987). This response strongly affects agronomic performance (yield, visual quality in ornemental plants; Borrell *et al*., 2014; Boumaza *et al*., 2010; Crespel *et al*., 2020; Slafer *et al*., 2014) and therefore must be finely tuned to growing conditions. Light in particular is a major regulator of branching and can vary strongly due to climatic fluctuations (cloudy periods) or artificial lighting in indoor production systems (Noia *et al*., 2023; SharathKumar *et al*., 2020). While several studies have investigated light quality (Demotes-Mainard *et al*., 2016), the impact of light quantity on bud outgrowth has received less attention. Most studies indicate that low light quantity represses branching (Corot *et al*., 2017; Demotes-Mainard *et al*., 2013b; Leduc *et al*., 2014; Mitchell, 1953a; Su *et al*., 2011), although transient low-light periods can stimulate branching (Demotes-Mainard *et al*., 2013b; Lafarge *et al*., 2010), highlighting the complexity of this response. The underlying physiological mechanisms remain debated (Schneider *et al*., 2019).

Historically, research has focused on apical dominance, the phenomenon by which growing apical organs inhibit the buds located below (Cline, 1991; Thimann and Skoog, 1933). Auxin, produced by apical organs and transported basipetally, inhibits bud outgrowth through interactions with cytokinins (CK) and strigolactones (SL) (Domagalska and Leyser, 2011; Schneider *et al*., 2019). However, growing evidence over the last decade has highlighted a major role of low sugar availability for buds, due to the strong sink activity of apical organs. A pioneering study in pea (Mason *et al*., 2014) showed that, following decapitation, basal bud release preceded any detectable decrease in auxin in adjacent stem tissues, whereas an increase in sugar levels was observed. The causal link between increased sugar availability and bud release was demonstrated by exogenous sugar supply in plants with intact apices, which promoted bud outgrowth.

Additional support for a key role of sugar availability comes from experiments manipulating sugar supply or demand for growth in various species. Partial defoliation reduced bud growth in sorghum (Kebrom and Mullet, 2015), while *tin* wheat mutants with earlier stem growth exhibited reduced branching (Kebrom *et al*., 2012). *In vitro* studies using isolated stem segments further demonstrated that increasing sugar concentration in the medium promotes bud outgrowth dose-dependently by enhancing both its frequency and the time at which bud outgrowth start (Barbier *et al*., 2015b; Henry *et al*., 2011; rose). For rose, experiments using non-metabolizable sucrose analogs showed that sucrose partly acts as a signalling molecule, interacting with hormonal pathways (Barbier *et al*., 2015b; Bertheloot *et al*., 2020; Rabot *et al*., 2012). Consistently, in pea, the level of trehalose-6-phosphate (T6P) in the bud, a proxy for sucrose availability, strongly correlated with bud outgrowth triggered by plant decapitation or exogenous sucrose supply *in vitro* (Fichtner *et al*., 2017). Increased sugar availability rapidly activates bud metabolism, making the bud an active sink for sugars (Henry *et al*., 2011; Wang *et al*., 2022; Wang *et al*., 2021).

Because light intensity strongly modulates photosynthesis and sugar production (e.g., Hirose and Werger, 1987), it has been hypothesized that light limitation inhibits bud outgrowth by reducing sugar supply, thereby reinforcing apical dominance. This view is indirectly supported by studies showing that the promoting effect of light on grass tillering is reduced at higher temperatures, which enhances organ growth and sugar consumption (Bos and Neuteboom, 1998; Mitchell, 1953b). More quantitative studies also reported correlations between tillering level and indices integrating radiation, leaf size and expansion rate at the critical period for tiller appearance (Alam *et al*., 2014; Alam *et al*., 2017; Kim *et al*., 2010). Accordingly, several ecophysiological models implement branching control via a competition index reflecting the balance between carbohydrate (C) production by source organs and C use by sinks for growth (Evers *et al*., 2010; Luquet *et al*., 2006; Mathieu *et al*., 2009; Warren-Wilson, 1972). While such index has been shown to reproduce sugar reserves dynamics under a specific condition, supporting that it is a good proxy for sugar availability, direct physiological evidence that sugar availability drives branching response to light is lacking.

In rose, sugar availability does not appear to be the limiting factor for bud outgrowth inhibition under low light, despite reduced stem sugar concentrations (Corot *et al*., 2017; Roman *et al*., 2016b). Exogenous sugar supply failed to stimulate bud outgrowth under low light, both in intact plants and in decapitated and defoliated plants kept in darkness. In contrast, exogenous application of CK, that displayed reduced levels under low light, restored bud outgrowth, indicating that CK rather than sugars are limiting. Similarly, increased tillering in wheat was observed in response to higher day-time light intensity, regardless of sugar concentration (Lorenzo *et al*., 2015). These results call for a reassessment of the role of sugars in the light intensity-mediated bud outgrowth.

Furthermore, the concept that light intensity primarily promotes bud outgrowth by increasing photosynthesis and carbohydrate supply needs to be reexamined, given that light also induces morphological and physiological adaptations that may counterbalance its effect on photosynthesis. Low light tends to stimulate organ elongation (e.g. internodes and leaves), enhancing light interception (Chenu *et al*., 2005; Dong, 1995; Garcia-Martinez and Gil, 2001; Hitz *et al*., 2019; Jiang *et al*., 2021), which may partially compensate for reduced light availability. Leaf appearance rate and expansion have also been reported to decrease (Baumont *et al*., 2019; Chenu *et al*., 2005; Gautier *et al*., 1999; Milthorpe and Newton, 1963), thereby reducing sink demand. Conversely, the photosynthetic apparatus acclimates to low light, decreasing photosynthetic capacity which may strengthen the impact of low light (Grindlay, 1997; Sims and Pearcy, 1992). Together, this highlights the need for a quantitative approach—currently lacking—to analyze how light intensity affects carbohydrate availability and bud outgrowth. Such an approach must also be dynamic, given the reported irreversible effects of transient low light on plant development and physiology (Granier and Tardieu, 1999a; Sims and Pearcy, 1992).

The objective of this study is therefore to re-evaluate the prevailing view that branching responses to light intensity primarily result from changes in sugar availability within the plant, using rose as model species. We investigated the effects of both continuous and transient light limitation, the latter leading to a counter-intuitive stimulation of branching (Demotes-Mainard *et al*., 2013b). Beside measuring bud outgrowth and sugar levels, our approach integrates detailed measurements of plant development and photosynthetic capacity into a computational model to dynamically quantify source-sink balance. Complemented by targeted source-sink manipulations, this integrative framework allows us to disentangle and explain the relationships between light intensity, sugar availability, and bud outgrowth.

## Material and methods

### Plant material, growth conditions, and light treatments

Single-node cuttings of *R. hybrida* ‘Radrazz’ plants were obtained as described in Demotes-Mainard et al. (2013). They were grown in 500 ml pots containing a 50/40/10 mixture of neutral peat, coconut fibers and perlite in a temperature-controlled greenhouse till 3 leaves were visible, and were kept disease-free. Then plants were transferred to a growth chamber (light/dark 16/8h photoperiod; 22/20°C day/night temperature; air humidity 60-70%) until the flower of the primary axis faded. Plants were sub-irrigated every two or three days with a fertilized solution (5.0 mM KNO3, 2.0 mM Ca(NO3)2, 2.0 mM NH4NO3, 2.0 mM KH2PO4, 2.0 mM MgSO4, 0.25 mM NaOH) and additional trace elements: Kanieltra formula 6-Fe at 0.25 ml.l^-1^ (Hydro Azote, Nanterre, France) to maintain optimal hydric and mineral nutrition.

In growth chambers, plants were submitted to three different light intensity regimes described in Fig. 1: a constant low light intensity (LL), a constant high light intensity (HH), or a transient light limitation until the floral bud becomes visible at the top of the primary axis (FBV stage; LH) followed by a return to high light intensity. A first experiment (Exp. 1) was made in 2011 in two identical growth chambers (Froids et Mesures, Beaucouzé, France), with a photosynthetic flux density (PPFD) of *ca*. 420 µmol m^-2^ s^-1^ at plant height (60 cm above the culture table) for high light, and *ca*. 90 µmol m^-2^ s^-1^ for low light. Plants were grown in 6 rows (including two border rows) with one plant every 15 cm. Since different growth chambers were used for the different light treatments, temperature was recorded, and time was expressed in degree days (2.1°C as base temperature; see Demotes-Mainard et al., 2013) to account for temperature differences between growth chambers. Other experiments (Exp. 2-3-4, 2018-2021) were made in a unique Strader growth chamber, with PPFD of *ca*. 80 µmol m^-2^ s^-1^ for low light and *ca*. 310 µmol m^-2^ s^-1^ for high light. Plants were grown in 2 rows with, similarly to Exp. 1, one plant every 15 cm. Since light treatments were applied concomitantly in the same growth chamber under a similar temperature, time was expressed in days and not degree days.

**Figure 1.**
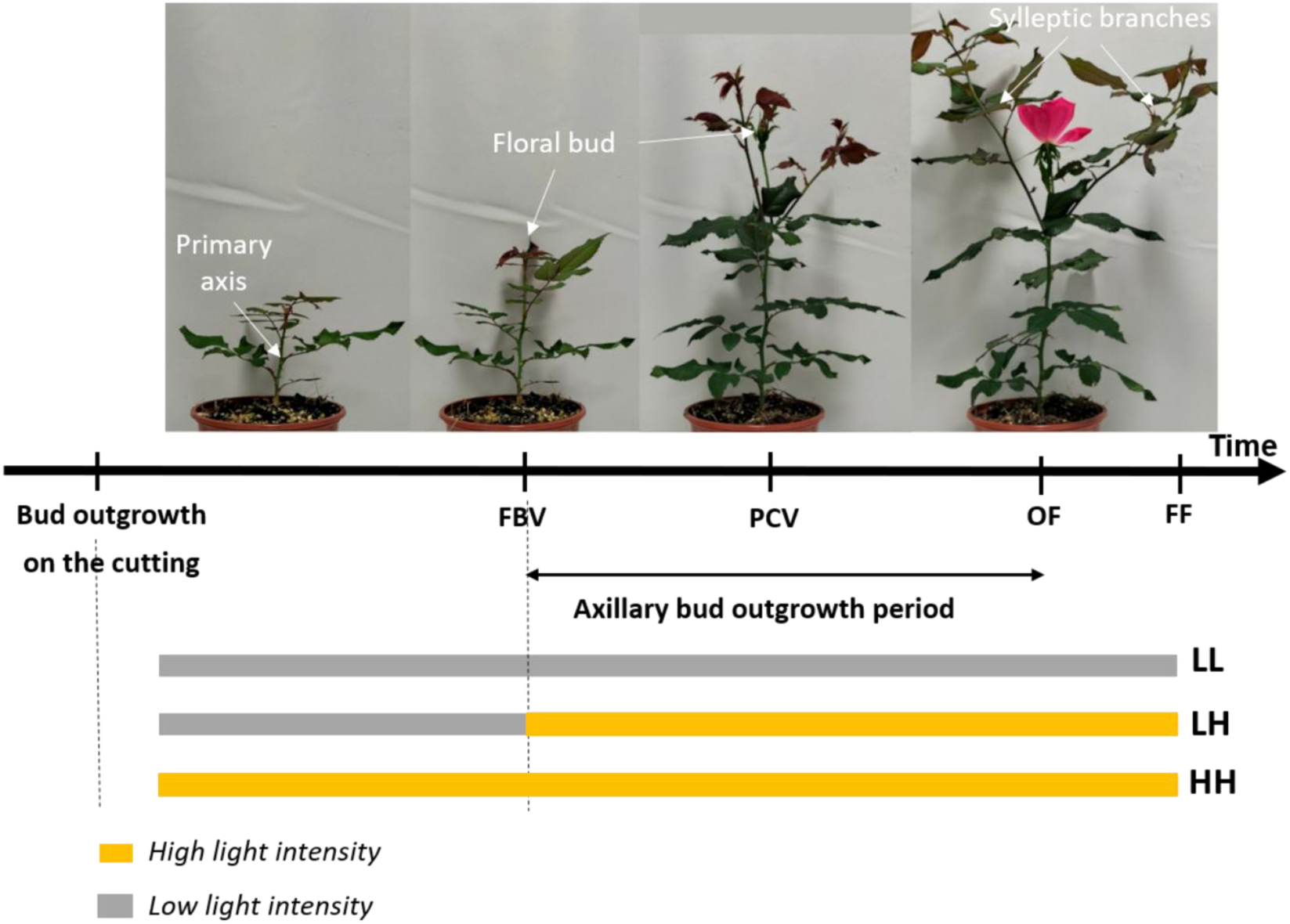
Light treatments applied to rose plants during their development. From plant emergence from a single-node cutting, plants were rapidly transferred to a growth chamber where they were submitted to different regimes of light intensity: (i) LL (Low-Low, grey strip): a constant low light intensity; (ii) HH (High-High, orange strip): a constant high light intensity; (iii) LH (Low-High): a temporary light limitation until the floral bud becomes visible at the top of the primary axis (FBV stage), i.e. before axillary buds started to grow out. *FBV:* floral bud visible; *PCV*: petal color visible; *OF*: opened flower: *FF*: faded flower.

### Plant architectural description

Studied rose plants are composed of primary axes that develop from the initial cutting (see Fig. 1). Each primary axis consists of a succession of vegetative phytomers, topped by a peduncle, that appear successively till the floral bud becomes visible at the top (FBV stage). Each phytomer consists of an internode, a leaf at the top, and an axillary bud at leaf axil, except for the most basal phytomers that are unleafy (called “basal part”). From the axillary buds, secondary axes develop after FBV stage. Each leafy phytomer of the primary axis was numbered from plant base, *n* being the total number of phytomers and *n*+1 being the peduncle. Phytomers were allocated to a zone, Z1, Z2, or Z3 (Fig. 2B; Fig. S2) according to their relative rank *r* (Z1: 0.7 ≤ *r* ≤ 1; Z2: 0.4 ≤ *r* < 0.7; Z3: 0 ≤ *r* < 0.4):

**Figure 2.**
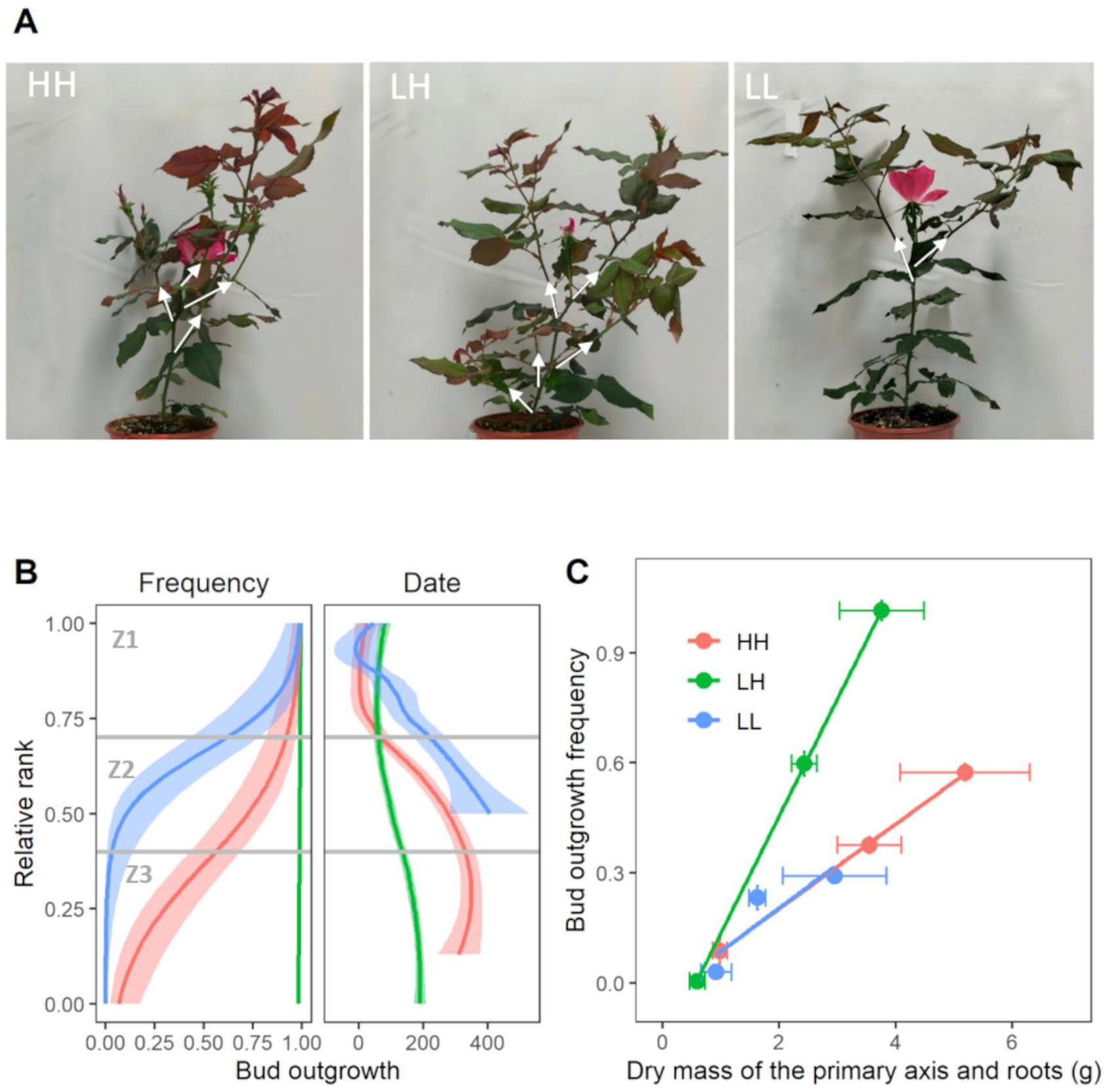
Axillary bud outgrowth was repressed by continuous light limitation but stimulated by transient light limitation compared to continuous high light. (***A***) Pictures of a rose plant at opened or faded flower stage (*OF*). White arrows represent lateral branches. (***B***) Outgrowth frequency and date (expressed as degree days after FBV) for buds at different ranks along the primary axis of rose plants grown under HH (red), LH (green), and LL (blue), in Exp. 1. Rank is expressed relative to the total number of leaves and assigned to a zone (Z1, Z2, Z3; see material and methods). For date, only ranks with a rate > 0.2 are represented. Lines represent logistic and loess smooth lines, respectively, ± 95% confidence intervals (*n*=15 plants for LH and 20 plants for HH, LL). (***C***) Relationship between (i) primary axis and root dry mass and (ii) bud outgrowth frequency for plants grown under HH, LH, and LL. Data are identical to Fig. S3 data. Symbols represent means for each variable, errors bars represent SEM, and lines the estimated linear regression between dry mass and bud outgrowth frequency for each light condition.

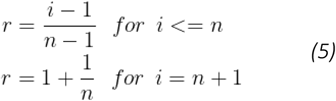

where *i* is the leafy phytomer number.

### Dynamic quantification of plant architectural development and growth (Exp. 1)

For Exp. 1, the rose canopy in each light treatment was divided in two groups: one group of 12 plants for non-destructive measurements, and one group of 24 plants for destructive measurements. For each primary axis of each group, the date at which the plant emerged on the cutting (bud outgrowth) and the floral stages (Fig. 1; FBV, floral visible stage; PCV, petal color visible; OF, opened flower; FF, faded flower) were recorded every one or two days.

In the first group, the number of visible leaves on each primary axis was counted every 1 or 2 days except for LH, which was assumed to follow a similar pattern compared to LL since leaves appear before FBV, so under the same PPFD for LL and LH. Leaf appearance date was estimated assuming a linear correlation between leaf relative rank and thermal time, as done earlier (Demotes-Mainard *et al*., 2013a). In addition, the development stage (dormant or outgrowing) of each axillary bud along primary axes was scored from FBV stage every 2 days. An axillary bud was considered to start its outgrowth when its first leaf was clearly visible between its scales.

In the second group, several sub-group of 4 to 5 plants were identified for harvesting at (i) FBV, just after plant transfer to high light for LH, (ii) *ca*. 130°Cd after FBV, and (iii) at or after the end of organ expansion. In the morning, each plant was separated into its root compartment, the primary axis basal part, each internode+node, each leaf, the peduncle, the floral organ, and each axillary branch. Then, each organ or compartment was immediately scanned (except roots and secondary branches), transferred in frozen nitrogen, stored at −80°C, lyophilized, and weighted. From scans, the area of each fully-opened leaf, the length of the terminal leaflet of each leaf, as well as the length and diameter of each internode was determined using ImageJ (https://imagej.nih.gov/ij/).

### Quantification of sugar contents (Exp. 1-2)

Samples of roots and primary axis individual organs or compartments of Exp. 1 were lyophilized then ground, and aliquots of 10 mg fractionated with H_2_O/EtOH as in Cross *et al*. (2006). Proteins were extracted by adding 400 µl NaOH 0.1 M to the pellet, vigorous shaking and heating at 95 °C for 30 min. After centrifugation at 2,000 g for 10 min, protein content was measured as in Bradford (1976). After neutralization with HCl 0.5 M, starch was digested and quantified in the the resuspended pellet as in Hendriks *et al*. (2003). Glucose, fructose and sucrose were measured in the supernatant as in Stitt *et al*. (1989), nitrate as in Tschoep *et al*. (2009) and total amino acids as in Bantan-Polak *et al*. (2001). All liquid handling steps were performed with Star and Starlet pipetting robots (Hamilton, Villebon sur Yvette, France). Extractions were performed in 1.1 ml Micronic tubes (Lelystad, The Netherlands) and all assays in polystyrene 96-well flat-bottom microplates. Absorbances were read at 600 (protein), 340 (starch and soluble sugars) and 540 (nitrate) nm in MP96 microplate readers (SAFAS, Monaco). Fluorescence (amino acids, 405 nm excitation, 485 nm emission) was measured in a Xenius fluorimeter (SAFAS, Monaco).

In addition, a complementary experiment, Exp. 2, was undertaken with the aim to specifically quantify metabolites in the primary axis before bud outgrowth period. Plants were grown in the same growth chamber under HH, LH, or LL, and four groups of at least four plants were pooled at (i) FBV, just before plant transfer to high light for LH, (ii) 4 days after FBV for HH and LH, and (iii) 7 days after FBV for HH, LH, LL. In each group, leaves and nodes (segments with 5 mm of stem each side of the node) of each zone of primary axes were separated and immediately frozen in liquid nitrogen and stored at −80°C in the morning. Samples were then lyophilized, ground, and soluble sugars (sucrose, glucose, fructose), starch, nitrate, total amino-acids, and soluble proteins and starch were quantified with the same method as in Exp. 1. In addition, total polyphenols were measured in the supernatant as in Colombié *et al*. (2023). In parallel, bud outgrowth was quantified as for Exp. 1, every 2 days from FBV, on a subset of 15-20 plants.

### *In vitro* culture of isolated stem segments

To test if an increase in sugar availability for buds is correlated to an increase in stem sucrose and starch contents (Fig. S4), a stem segment with one bud was excised from 10 rose primary axes at FBV stage, and grown two days on a medium supplemented with either mannitol (100 mM) or sucrose at different concentrations (10, 50, 100, 250 mM), as described previously (Barbier *et al*., 2015b). 48h after transfer on the culture medium, stem segments were collected in liquid nitrogen, and sugar content analyses were performed on three independent biological samples using 50 mg of frozen tissues per sample, similarly to Bertheloot *et al*. (2020).

### Quantification of photosynthesis variables and final dimensions of primary axis organs (Exp. 3)

Fourteen to seventeen plants were grown under HH, LH, or LL, and for each plant, the date at which the plant emerged on the cutting and the number of visible leaves were recorded each day. When the fourth leaf was visible, 3 times per week, the length of the terminal leaflet of each leaf was measured, the floral stage of the primary axis recorded, and the chlorophyll index of each leaflet measured using Dualex device (Dualex 4 Scientific, Cerovic *et al*., 2012) till OF stage. Chlorophyll index of each leaf was calculated as the mean of the measurements per leaflet. In addition, net photosynthesis was measured on a subset of 5 plants at 3-4 days after FBV for the terminal leaflet of leaves at ranks 1 and 4, at 100 and 600 μmol m^-2^ s^-1^ PPFD, using LICOR device (LI-6400 XT, LI-COR, Lincoln, NE, United States; 10/90% blue/red light, 400 μmol CO_2_, 70% relative humidity, 22°C for temperature). On another subset of 10 plants, PPFD at each leaf level was measured at 3, 10 and 14 days after FBV using a quantum sensor (LI-190R Quantum Sensor, LI-COR, Lincoln, NE, United States) placed near the terminal leaflet and pointing upward.

At OF stage -once primary axes had finished their development-, each leaf and internode of each plant under HH and LH was separated similarly to Exp. 1, scanned, and leaf areas, terminal leaflet lengths, and internodes/peduncle lengths and diameters were determined using ImageJ (https://imagej.nih.gov/ij/).

### Estimation of plant C availability

C availability *I*_*c*_ was estimated at each time step *t* (°Cd) for a primary axis of a rose plant grown in Exp. 1 under HH, LH, or LL, from FBV stage till *ca*. 300 °Cd after FBV (period of sugar accumulation for all treatments).

#### *I*_*c*_ calculation model

*I*_*c*_ (dimensionless) was calculated as the ratio between C supply (mg C °Cd^-1^) by photosynthesis and C demand (mg C °Cd^-1^), defined as C needed to build the structure of plant organs (Luquet *et al*., 2006; Tabourel-Tayot and Gastal, 1998):

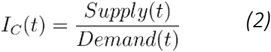

In this way, an *I*_*c*_ above 1 indicates an excess in C while an *I*_*c*_ below 1 indicates a C shortage. Both the supply and demand were dependent on the temporal expansion dynamics of plant organs. The increases in leaf *i* area, *A*_*i,lf*_ (cm^2^), and internode *i* length, *L*_*i,m*_ (cm), including the peduncle, were described by logistic functions of time saturating at maxima 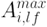 (cm^2^) and 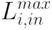 (cm) respectively, with normalized maximum rates, *w*_*lf*_ and *w*_*in*_ (°Cd^-1^), at 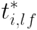 and 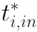 (°Cd), respectively (Demotes-Mainard *et al*., 2013a):

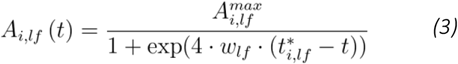

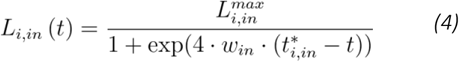

The supply was the sum of each leaf *i* photosynthesis, calculated as the product of leaf *i* area and surfacic photosynthesis, *P*_*i,lf*_ (μmol C m^-2^ s^-1^), that was estimated as a rectangular hyperbolar of intercepted PPFD (Equatiion 9), *PPFD*_*i,lf*_ (μmol photons m^-2^ s^-1^) (Thornley, 1998):

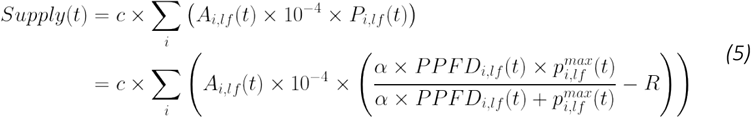

where *c* is a conversion coefficient (mg C μmol C^-1^ s °Cd^-1^), *R* is the respiration (assumed constant, μmol m^-2^ s^-1^), *α* the photosynthetic efficiency (mol C. mol photons^-1^), and 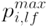 (μmol C m^-2^ s^-1^) the maximum photosynthesis at saturating PPFD level depending on leaf rank and its development stage as detailed below.

C demand is the C used at each time step to build the structure of plant organs:

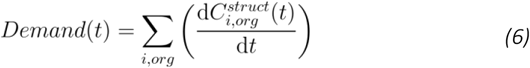

with 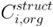 being the C involved in organ (*org*) *i* structure, organ being the leaves, the internodes, the basal part, the flower of the primary axis, as well as roots and secondary axes.

For the leaves and internodes, the C demand was decomposed as C used for the increase in dimensions (leaf area, internode volume expressed as a function of length and diameter *D*_*i,in*_ (cm)) and for the increase in thickness (structural C mass per unit area, *CA*_*i,lf*_ (mg C cm^-2^), for leaves, and unit volume, *Cv*_*i,in*_ (mg C cm^-3^), for internodes).

For the leaves, the C demand is thus:

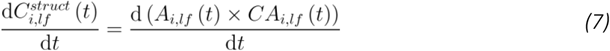

And for the internodes, it is formalized as:

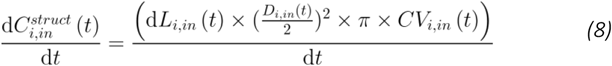

#### *I*_*c*_ estimation

From measurements of leaf appearance, final organ dimensions, photosynthesis, chlorophyll index, and organ dry masses, *I*_*c*_ was estimated each 10°Cd for median plants that were grown under HH, LH, or LL in Exp. 1 using equations above. The primary axis of these median plants was made of 8 leafy phytomers for LL and LH, and 9 for HH (Fig. S6). In measurements, axes may have a different number of phytomers compared to the median plant in each treatment. Estimations were made after attributing to each measured organ at rank *i* a median plant-equivalent rank, calculated from the actual relative rank and the targeted median number of phytomers, according to equation 1.

#### Estimations of leaf and internode expansions (Equations 3-4)

On the one hand, inflexion dates of individual leaves and internodes, 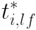 and 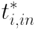 respectively, were estimated as pre-established polynomial functions of measured leaf appearance dates for each individual plant in Exp. 1 (Demotes-Mainard *et al*., 2013a). Then, a cubic smooth spline was adjusted on inflexion dates as a function of the relative rank, for HH and LL, giving access to the average inflexion dates for individual leaves and internodes of a median primary axis in HH and LL (Fig. S9A). Estimated inflexion dates of LL were used for LH, since LL and LH organs appeared under similar low light intensity. On the other hand, maximum areas of individual leaves 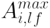 and maximum lengths of individual internodes 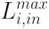 were estimated as means of areas and lengths, respectively, per median plant-equivalent rank (Fig. S9B). After statistical analysis on measured data, similar values were set up for leaf areas for LH and HH for ranks ≤ 5 as well as LL and HH for rank > 5. For internodes, similar values were set up for all treatments for ranks ≤ 5. Finally, by integrating estimated inflexion point dates and final dimensions in equations 3-4, *w*_*lf*_ and *w*_*in*_ were manually adjusted so that estimated leaf and internode expansion dynamics, *A*_*i,lf*_ and *L*_*i,in*_, fitted the measured ones (Fig. S10-11).

#### Supply estimation (Equation 5)

In Exp. 1, *PPFD*_*i,lf*_ was approximated as a function of cumulative leaf area index (*LAI*, leaf area per unit soil area) above leaf *i* and incident PFFD (*PPFD*_0_), using the Beer-Lambert’s law:

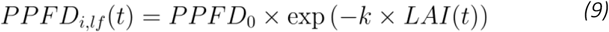

where *k* is the light extinction coefficient. *k* was estimated using Exp. 3 data of PPFD (Fig. S7) and corresponding estimated leaf areas. Leaf expansions were estimated with the methodology described for Exp. 1 from measured leaf appearance dates and maximum areas (data not shown).

To estimate the surfacic photosynthesis *P*_*i,lf*_ response to *PPFD*_*i,lf*_, the parameters of the photosynthesis response curve to PPFD, *α* and 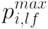, were first estimated on Exp. 3 data by fitting (Equation 5) to photosynthesis measured for ranks 1 and 4 at 100 and 600 μmol m^-2^ s^-1^ PPFD (Fig. 4A). Then, a linear dependence of 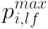 with leaf N status (Bertheloot *et al*., 2011; Grindlay, 1997) depending on leaf age and rank, was introduced using leaf chlorophyll index, *chl*_*i,lf*_, as a proxy for leaf N status (Bertheloot *et al*., 2011). Temporal dynamics were estimated from Exp. 3 chlorophyll measurements after statistical analysis (ANOVA) to determine if time-, rank-, treatment-effects had to be accounted for: *chl*_*i,lf*_ started with similar values for all treatments and leaves at the time of leaf appearance and linearly increased till maximum chlorophyll content was reached. The increase rate was higher in HH than LL, and after FBV than before FBV for LH leaves expanding after FBV (Fig. S15). These functions describing chlorophyll temporal dynamics were applied to estimate chlorophyll dynamics in Exp. 1 from the leaf appearance dates in this treatment. 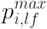 was set proportional to estimated *chl*_*i,lf*_ (proportion coefficient estimated from Exp. 3 data).

Once estimated, the surfacic photosynthesis of each leaf, *P*_*i,lf*_, was multiplied by leaf area, and each leaf photosynthesis summed, giving the total photosynthesis of the axis (Equation 5). C supply was obtained by multiplying photosynthesis by a conversion coefficient *c* that accounts for a 16h-photoperiod and a mean 20.6 °Cd per day according to Exp. 1 temperature measurements.

#### Demand estimation (Equations 6-8)

Firstly, 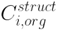 of each organ or compartment of each plant in Exp. 1 at each date was calculated as the difference between the measured total C mass (assuming 0.45 gC. gM^-1^, M being dry mass) and the measured C mass allocated to soluble compounds (hexoses, sucrose, starch, amino acids, soluble proteins, and polyphenols). 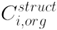 for each leaf and for each internode was then divided by the measured leaf area or internode volume, respectively, to get the measured *CA*_*i,lf*_ and *CV*_*i,in*_ Secondly, a linear fitting procedure was applied on the measured 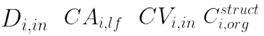 (*org=* base, roots, flower) to obtain the temporal variation of these variables (Fig. S12-S14). This was preceded by a statistical analysis (ANOVA) to determine if time-, rank-, treatment-effects had to be accounted for. In all cases, similar initial values were taken for all organs and treatments when data were unsufficient to estimate these values. Finally, the demand was calculated for each organ/compartment each 10°Cd, using equations 7 and 8 for individual leaves and internodes, and directly from the temporal variation of the other compartments.

### Manipulation of source-sink balance (Exp. 4)

Sugar source for plants in HH and LL was artificially increased by exogenous sugar supply at FBV in zone Z3 where differences in bud outgrowth were strongly marked between light treatments. The terminal leaflet of the higher foliated leaf of zone Z3 was removed and the corresponding petiole was immersed in a aqueous solution (with 0.25% of Preservative Plant Mixture) of either 100mM sucrose or 50mM mannitol, an osmotic control, as made in Bertheloot et al. (2020).To avoid the diversion of sugar supplied in this way towards the actively growing apical organs, additional sucrose (100 mM) was vascularly supplied above Z3 (through the lowest internode of Z2), using the cotton wick-method described in Corot et al. (2017), for both petiole-fed sucrose and mannitol treatments (Fig. S1). This was done with the intention to feed the growing apical organs through a supply in Z2 and enable testing the effect of an increase in sugar locally in Z3.

Sugar source for plants in LH was artificially decreased by either masking the leaves or by applying a photosynthesis inhibitor. In the masking experiment, the upper surface of leaflets of all fully opened leaves were covered using either a black opaque (treated) or a transparent plastic film (control) from the fifth days after FBV. In the photosynthesis inhibition experiment, a solution containing DCMU (DCMU 50µM, 0.5% ethanol, Tween) or a similar solution without DCMU was applied once with a brush on half of each fully opened leaf from the sixth day after FBV. In addition, at FBV+7 days, photosynthesis was measured on leaf 4 (median leaflet) of treated and non-treated plants with DCMU to verify that photosynthesis was indeed decreased by DCMU application (data not shown).

For each treatment, the stage of development of each bud was recorded every two or three days.

### Statistics and fitting procedures

R software for Windows was used for statistical analyses, fitting procedures, and supply/demand calculation model.

The functions aov(), TukeyHSD(), fisher.test(), t.test() were used for analysis of variance (ANOVA), Tukey multiple comparison test, Fisher’s exact test, Student’s test, respectively. The functions geom_smooth(method=‘lm’), geom_smooth(), and geom_smooth(method = glm, method.args= list(family=“binomial”)) to estimate linear regressions, trends lines for continuous variables (*e*.*g*. time of bud outgrowth, chlorophyll) and for qualitative variable (bud outgrowth frequency), respectively. The parameter estimations of specific functions (describing the temporal dynamics of chlorophyll, diameter, structural mass per unit area or volume, or PPFD attenuation with LAI, photosynthesis response to PPFD) was done by minimizing the root mean square errors between observations and estimations using the nlm() procedure.

## Results

### Continuous light limitation repressed axillary bud outgrowth, whereas transient light limitation overstimulated it and increased plant investment in bud outgrowth relative to primary axis growth

In Experiment 1, single-axis rose plants derived from cuttings were grown under continuously high (*ca*. 420 μmol m^-2^ s^-1^, HH), continuously low (*ca*. 90 μmol m^-2^ s^-1^, LL), or transiently low light (Low-High: LH) with a shift to high light applied just before the onset of bud outgrowth at the floral bud visible stage (FBV stage; Fig. 1). Each bud developmental status (dormant vs. outgrowing) and outgrowth timing (referred to as “outgrowth date”) were recorded along the primary axis. The axis was divided into an apical zone (Z1; mainly including buds that do not enter dormancy), an intermediate zone (Z2), and a basal zone (Z3; Fig. 2B).

Light treatments strongly affected bud outgrowth and branching (Fig. 2A-B). Under HH, bud outgrowth (Fig. 2B) and branch development (Fig. 2A) were mainly restricted to Z1 and Z2 and followed a top-to-bottom sequence in outgrowth dates (Fig. 2B). Under LL, this pattern was preserved but bud outgrowth frequency was reduced in Z2 and Z3 (7% *vs*. 44% in HH on the entire axis). In contrast, LH induced branching along the entire axis, with 98% of buds growing out in Z2 and Z3. Bud outgrowth was more synchronous, resulting in a significant earlier mean bud outgrowth date all over the axis (146, 254, and 356 °Cd for LH, HH, and LL, respectively). These effects were confirmed in a second experiment performed n a different growth chamber (Exp. 2; Fig. S2).

Because light is known to affect plant development (e.g., Chenu *et al*., 2005; Gautier *et al*., 1999), we examined whether differences in bud outgrowth between light treatments were associated with differences in primary axis growth. Dry mass of the different organs of the primary axis and roots was quantified at several time points after FBV stage (floral bud visible at the axis top), and compared with the bud outgrowth frequency cumulated along the primary axis (Fig. S2). Bud outgrowth frequency was strongly related to dry mass accumulation for all light conditions, but LH displayed a stronger bud outgrowth frequency increase relative to dry mass accumulation (Fig. 2C) indicating a higher relative carbon allocation to axillary bud outgrowth under transient light limitation.

### Continuous light limitation decreased both starch and sucrose contents, while transient light limitation promoted starch over-accumulation

To assess the relationship between light-dependent bud outgrowth patterns and sugar availability, sucrose and starch were quantified in stems (internodes+nodes) and leaves of the median (Z2) and basal (Z3) zones, which displayed clear differences in bud outgrowth accross treatments (Fig. 2B). Sucrose and starch were used as indicators of sugar availability (Kool *et al*., 1996), consistent with *in vitro* rose stem segments experiments showing dose-dependent increases in starch and sucrose concentrations with increasing external sucrose supply (Fig. S4; Barbier *et al*., 2015b).

In experiment 1, samples were collected at three time points: (1) FBV (immediately after the light shift in LH), (2) *ca*. 130 °Cd after FBV, corresponding to the onset of bud outgrowth in HH and LH, and (3) after the end of bud outgrowth period for HH (*ca*. 400 °Cd) and LH (*ca*. 240 °Cd), and at its onset for LL (*ca*. 290 °Cd) (Fig. 3A).

**Figure 3.**
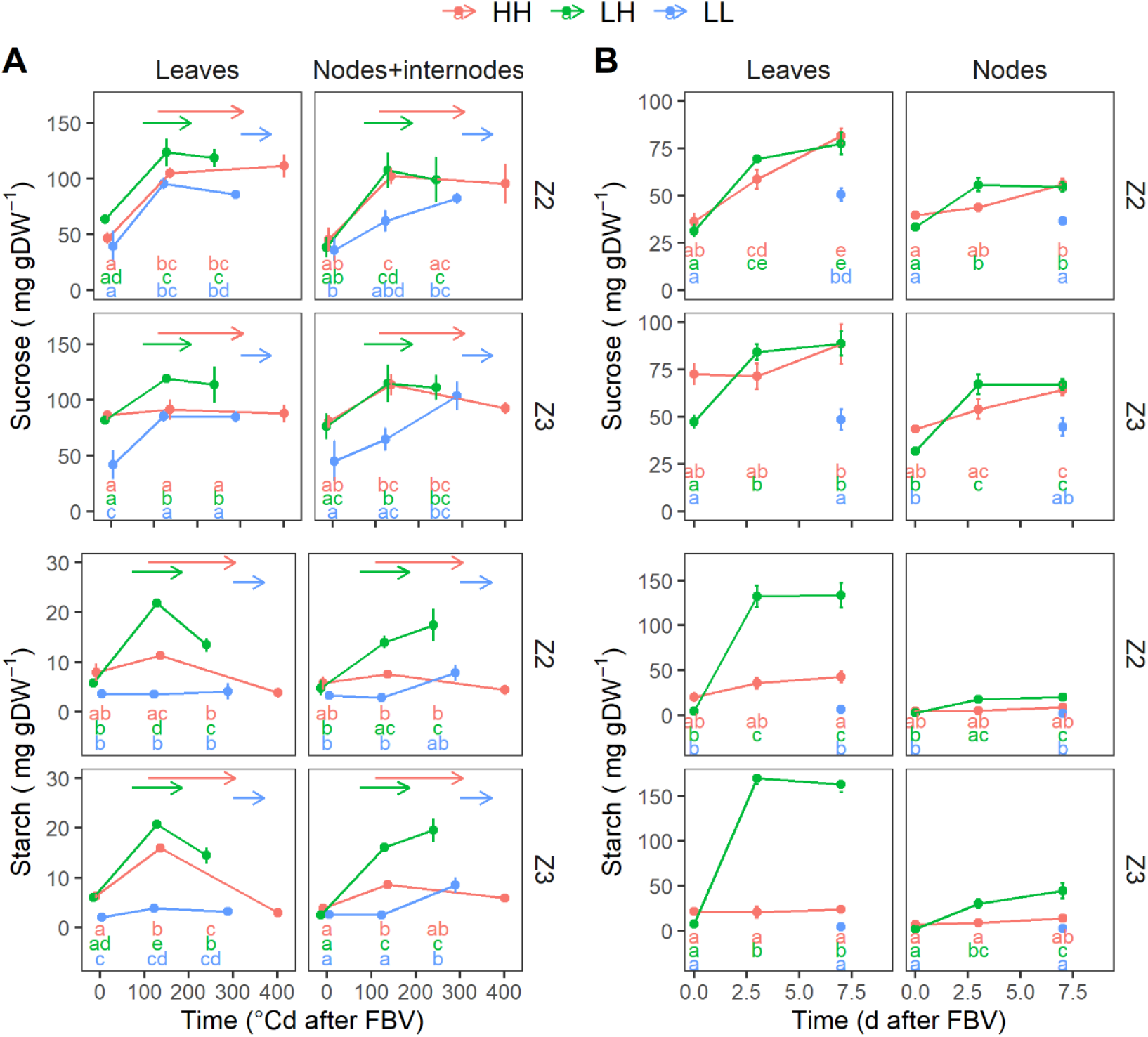
Sucrose and starch levels were reduced after FBV by continuous light limitation but stimulated by transient light limitation. Temporal dynamics of sucrose and starch for zones Z2 and Z3 of rose plants grown under HH (red), LH (green), or LL (blue) in Exp. 1 (***A***) and Exp. 2 (***B***). In (*A*), sugars were quantified in the leaves or internodes+nodes of each zone all over primary axis growth, including bud outgrowth period; horizontal arrows indicate bud outgrowth period, with the start of the arrow being the date at which bud outgrowth starts in Z2. Data are means ± SEM (*n*=4-5 plants). Time is expressed in degree days to account for the slight difference in temperature between the two growth chambers where low and high PPFD were applied, respectively. In (*B*), sugars were quantified in the leaves or nodes of each zone, specifically before the beginning of bud outgrowth. Data are means ± SEM of 4 repetitions of at least four pooled plants each. Time is expressed in days. For (*A,B*), different letters indicate significant differences between treatments and sampling dates (ANOVA followed by a Tukey test, *p* < 0.05). In (*B*), no measurement was made for LL plants at time=0 considering an identical value to LH plants (shifted to high light just after time 0; blue letter written but identical to the green letter).

**Figure 4.**
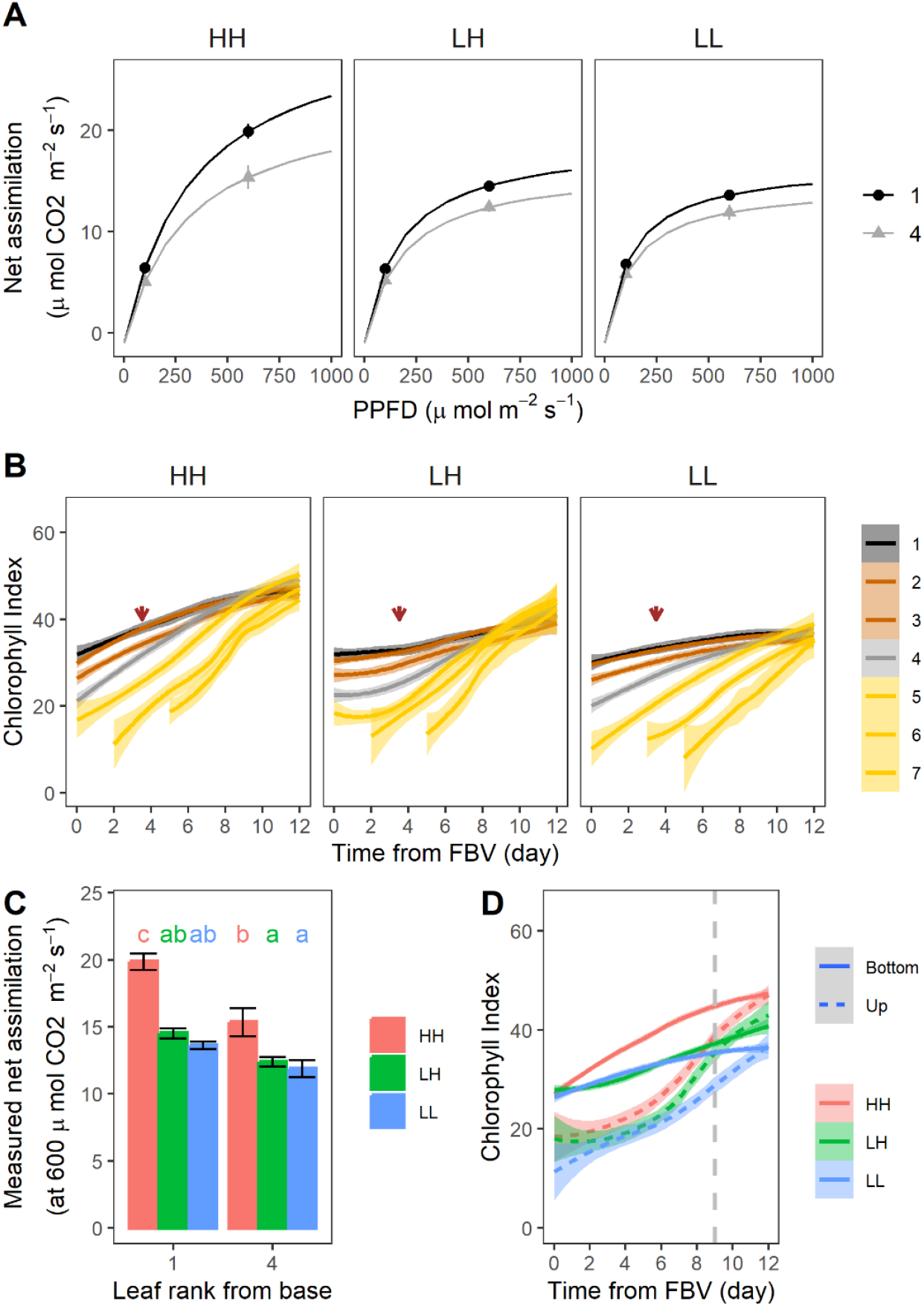
Leaf photosynthesis capacity was reduced after FBV for continuous and temporary light limitation. (***A***) Measured (symbols) and fitted (lines) CO_2_ net assimilation of mature leaves 1 (black) and 4 (grey), counting from base, of rose primary axes grown under HH, LH, or LL in Exp. 3. Measurements were made 3 or 4 days after FBV at a PPFD at leaf level of 100 or 600 μmol m^-2^ s^-1^, and are means ± SEM of 5 plants. Asterisks indicate significant differences between leaf ranks at each PPFD level (Student’s test, p<0.05). Estimations were made by fitting a rectangular hyperbola (Equation 5). Measured chlorophyll index as a function of time from FBV for each leaf rank (each line). Data represent loess regressions ± 95% confidence intervals (*n*=15-17 leaves). The brown vertical arrow represents the date at which net assimilation was measured in (*A*). (***C***) Focus on the measured net assimilation at 600 μmol m^-2^ s^-1^ of (A). Different letters indicate a significant difference between ranks and light treatments (pairwise comparison using Tukey test; *P*<0.05). (***D***) Overall chlorophyll index dynamics for the lower fully-expanded leaves (ranks <=4) and the upper expanding ones (ranks >4; *D*; Fig. S5). Data represent loess regressions ± 95% confidence intervals (*n*=45-68 leaves). The vertical dashed grey line represents the date at which bud outgrowth started.

Under HH, sucrose and starch contents increased between FBV and the onset of bud outgrowth (*ca*. 130 °Cd), significantly for sucrose in Z2 and starch in Z3. During the subsequent bud outgrowth period, they stabilized in stems and leaves, except for leaf starch that decrased. In LH, the increase in sucrose and starch between BFV and *ca*. 130 °Cd was significant for both Z2 and Z3, with markedly higher starch accumulation than in HH. Consequently, at the onset of bud outgrowth, LH plants had accumulated more starch than HH plants, after which sugar levels stabilized or declined similarly to HH. In LL, sucrose increased significantly only in leaves, while starch remained unchanged between FBV and *ca*. 130 °Cd in leaves and stems; both sugars accumulated later in stems, between 130 °Cd and the onset of bud outgrowth (*ca*. 290 °Cd). Overall, in all treatments, bud outgrowth started only after sucrose and starch had accumulated in stems, with delayed accumulation in LL and enhanced starch accumulation in LH.

The second experiment (Exp. 2; Fig. 3B), focusing on the period before the onset of bud outgrowth period in all treatments (*ca*. 8 days; Fig. S2), confirmed these patterns. Measurements were taken at 0, 4, and 7 days after FBV for HH and LH, and at 0 and 7 days for LL (using the same plants for LH and LL at time 0, prior to the LH light shift). Starch accumulated more in LH than in HH, whereas neither sucrose nor starch accumulated significantly in LL (except for sucrose in Z2, which remained lower than in HH at day 7).

Together, these results reveal a strong correlation between bud outgrowth, and the dynamics of sugar accumulation in stems, supporting a potential role for sugar availability in the light control of bud outgrowth.

### Both continuous and transient light limitation reduced the photosynthetic capacity of mature leaves after FBV

To determine whether differences in sugar availability between treatments were linked to photosynthesis, a third experiment (Exp. 3) quantified the photosynthetic capacity of primary-axis leaves under HH, LH, and LL (Fig. 4). Photosynthetic capacity reflects the ability of leaves to fix C and is closely associated with nitrogen content, as a major component of the photosynthetic machinery (Evans, 1989; Grindlay, 1997).

Net CO_2_ assimilation was first measured, 3-4 days after FBV, on two fully-expanded basal leaves (ranks 1 and 4) under low (100 μmol m^-2^ s^-1^) and high (600 μmol m^-2^ s^-1^) photosynthetic photon flux density (PPFD), and light response curves were fitted (Fig. 4A). Across treatments, older leaves (rank 1) exhibited higher photosynthetic capacity than younger ones (rank 4), consistent with age-dependent increases in nitrogen, RubisCo, and chlorophyll content during early leaf development (Osaki *et al*., 1995). HH plants displayed higher photosynthetic capacity than LH and LL plants (Fig. 4A, C), indicating a lasting acclimation to the lower light experienced during leaf expansion in LH (the four most basal leaves had expanded under low light), as previously reported (Oguchi *et al*., 2005; Sims and Pearcy, 1992).

To extend these observations to all leaves throughout their expansion, chlorophyll (Chl) index was used as a proxy for leaf nitrogen content (Evans, 1989). As expected, in fully-expanded lower leaves (ranks 1 to 4), Chl index increased with leaf age (Fig. 4B) and was lower in LH and LL compared to HH (Fig. 4B black and red, or Fig. 4D full lines). In upper expanding leaves, Chl index was lower than in mature leaves for all treatments, increased over time and was: (i) slightly lower in LH than in HH, (ii) the lowest in LL from day 7 onwards, *i*.*e*. before the onset of bud outgrowth (*ca*. 9 days). Thus, before bud outgrowth, whole-axis photosynthetic capacity was reduced in both LH and LL compared to HH, mainly due to lower photosynthetic capacity in mature basal leaves.

### Continuous light limitation reduced photosynthesis at the axis scale, but not transient light limitation

At the whole-plant scale, photosynthesis depends on the photosynthetic capacity of individual leaves, the PPFD they receive, and their area. Leaf area determines both the overall photosynthetic surface and PPFD attenuation from the top to the bottom of the canopy (Varletgrancher *et al*., 1989). We therefore calculated primary-axis photosynthesis during the sugar accumulation period in Exp. 1 (FBV to 300°Cd), using a model integrating these components (see Material and methods for calculation details).

Leaf expansion was estimated using an existing model for the same rose cultivar (Demotes-Mainard et al., 2013), fitted to our data. Lower mature leaves were the largest in LL, consistent with a shade avoidance response (Grime, 1981), whereas upper expanding leaves were the smallest in LH (Fig. 5A). HH plants also produced more leaves on average than LL and LH (Fig. S6).

**Figure 5.**
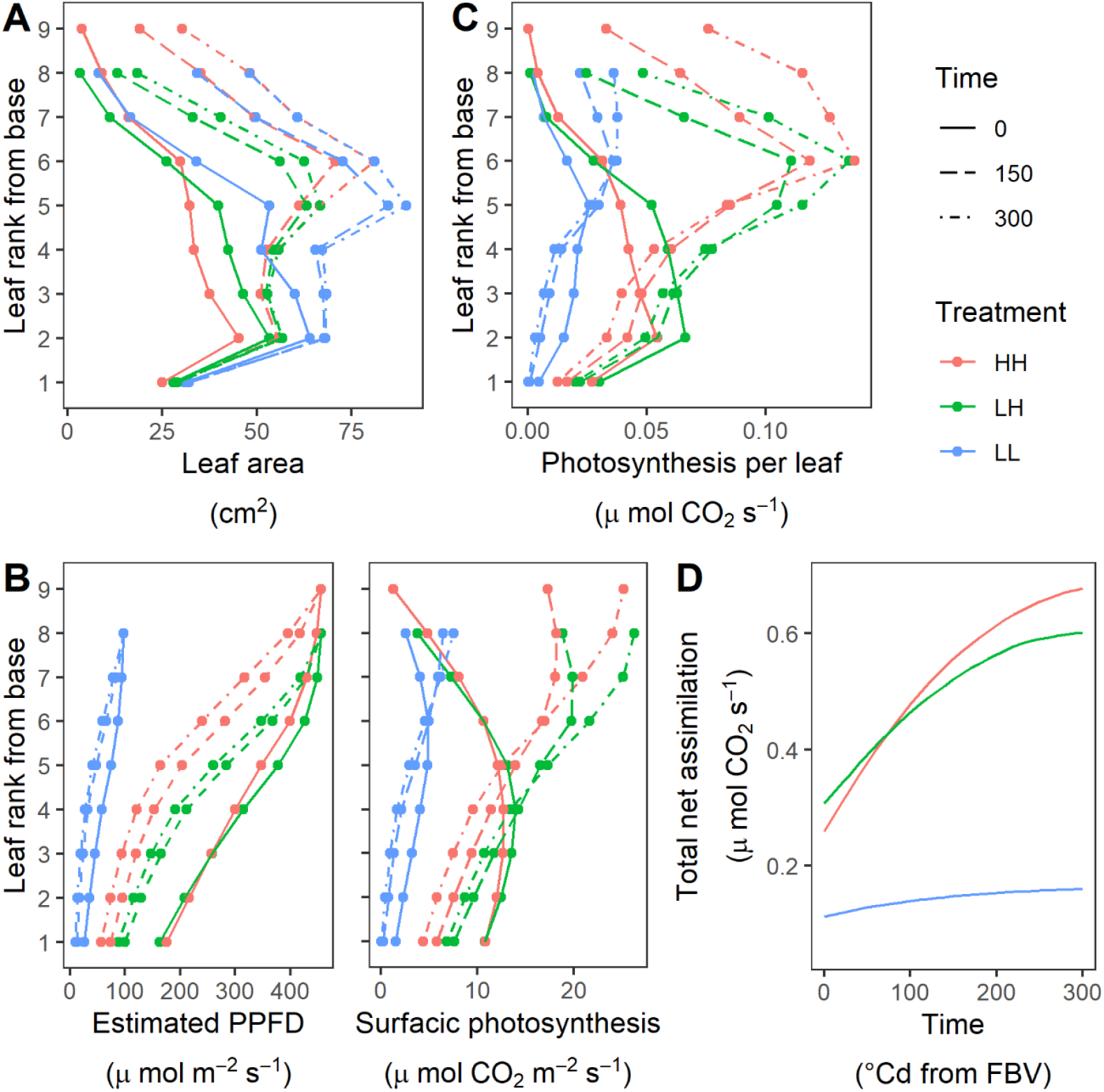
Estimated photosynthesis at primary axis scale was reduced by continuous but not temporary light limitation. Estimated temporal dynamics - between 0 and 300 °Cd - of the photosynthesis determinants and of photosynthesis of primary axes of rose plants grown under HH, LH, or LL in Exp. 1. (***A***) Leaf areas, (***B***) local PPFD and surfacic photosynthesis, (***C***) absolute photosynthesis of individual leaves along the primary axis at 0, 150, and 300 °Cd, and (***D***) photosynthesis of the primary axis through time. Methods for estimations are detailed in material and methods. The estimated primary axes had 9, 8, 8 leaves for HH, LH, LL, respectively, according to the median total number of leaves in each light treatment.

Each leaf was assigned a photosynthetic capacity estimated on Chl indices and a PPFD environment derived from the cumulated leaf area above it. As expected, PPFD was the lowest in LL due to low incident light (Fig. 5B left). In contrast, despite receiving the same incident light as HH, LH plants experienced higher PPFD at lower leaf levels owing to reduced shading from smaller upper leaf, a pattern confirmed by direct measurements (Fig. S7; Exp. 3) and previous light simulations on Exp. 1 plants reconstructed in 3D (Demotes-Mainard *et al*., 2025).

Using PPFD and photosynthetic capacity, surfacic photosynthesis was computed for each leaf (*P*_*i*_ in equation (5); Fig. 5B right). PPFD differences primarily drove treatments effects, resulting in the lowest surfacic photosynthesis in LL and the highest in LH. The reduced photosynthetic capacity in LH (Fig. 4) therefore had little impact on surfacic photosynthesis compared with HH. Photosynthetic capacity instead mainly affected surfacic photosynthesis during the period of leaf expansion, resulting in low initial values for upper leaves (at °Cday), followed by increases as leaves expanded (at 150 and 300°Cday).

When multiplied by leaf area, surfacic photosynthesis estimates yielded absolute leaf photosynthesis estimates (Fig. 5C). These were the lowest in LL, despite larger lower leaves, because of low PPFD. Comparing LH with HH, basal leaves showed higher photosynthesis due to reduced self-shading, while upper-leaf photosynthesis was lower because of smaller leaf size. Consequently, total primary-axis photosynthesis was similar in LH and HH, but 3-6 times lower in LL (Fig. 5D). It increased over time with leaf expansion for all treatments, although more slowly in LL due to PPFD limitation.

Thus, compared with HH, reduced sugar contents in LL can be explained by limited photosynthesis, whereas sugar over-accumulation in LH cannot be attributed to increased photosynthetic activity.

### Transient light limitation before FBV reduced organ expansion after FBV

As reported for water stress (Muller *et al*., 2011; Tardieu *et al*., 1999), the sugar over-accumulation in LH relative to HH could result from reduced organ expansion, lowering C demand while C supply from photosynthesis remains similar, thus increasing supply-demand ratio.

In addition to the smaller estimated upper-leaf areas in LH compared with HH (Fig. 5A), measured organ expansions between FBV and 130°Cd later (corresponding to the sugar accumulation period) in Exp. 1 supported this hypothesis (Fig. 6). Upper leaves (relative rank > 0.55)– that appeared under low light (Fig. S5)-expanded more in HH than in LL or LH (Fig. 6A), indicating that light conditions during early developmental stages had a lasting effect on subsequent expansion. This aligns with previous findings (Granier and Tardieu, 1999a) and with the reduced final area of upper leaves observed in LH compared with HH in a separate experiment (Exp. 3; Fig. S8A). Internode development was also affected by early light conditions: diameter was reduced in both LH and LL compared with HH (Fig. 6B; Fig. S8B) while length showed a weaker but non-significant reduction in Exp. 1 (Fig. 6C; Fig. S8C). Overall, shortly after FBV, both LH and LL plants exhibited reduced organ expansion compared with HH, supporting the hypothesis that reduced expansion contributed to sugar over-accumulation in LH.

**Figure 6.**
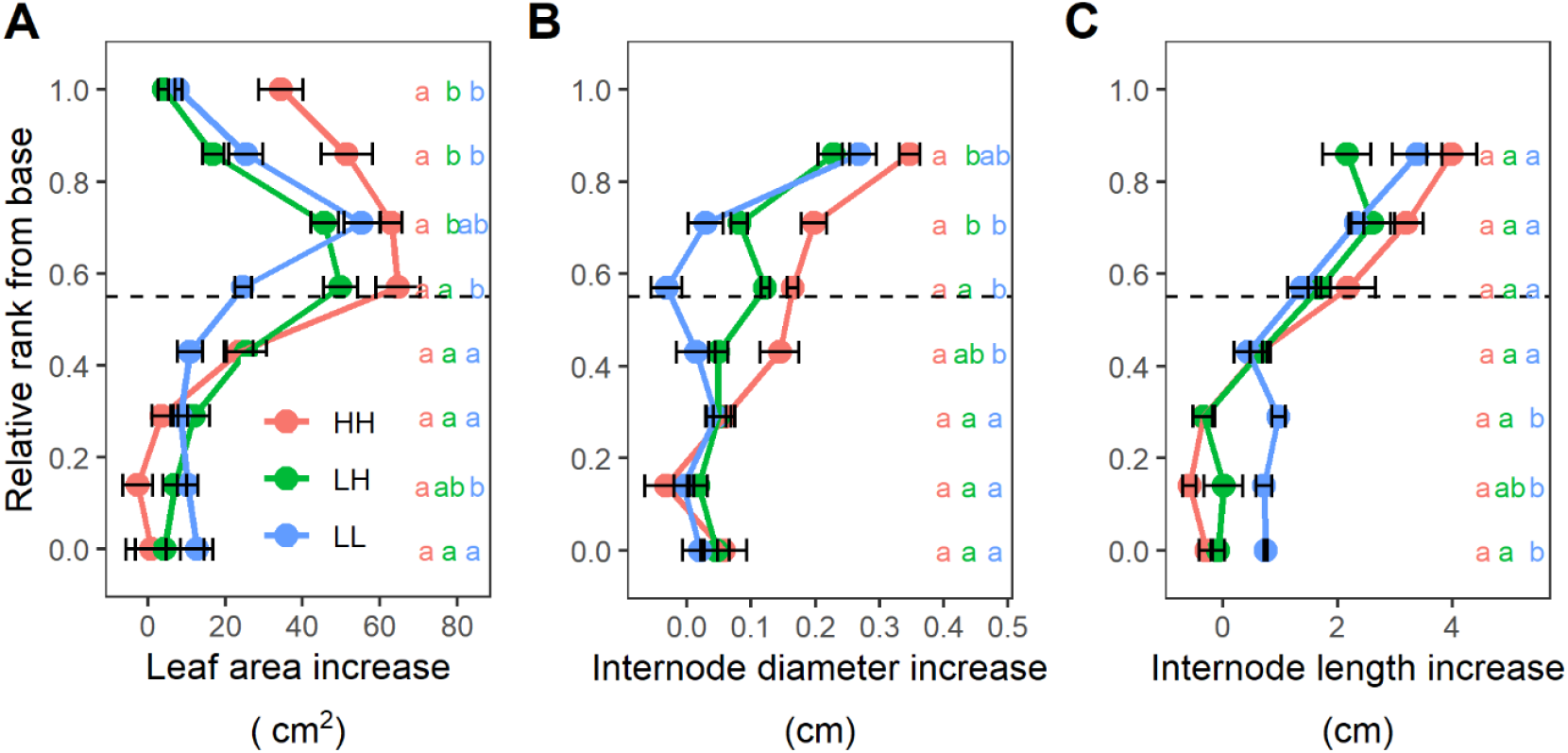
Low light, either continuously or transiently applied, reduced the increase in dimensions of primary axis organs after FBV. Increase - between FBV stage and FBV + *ca*. 130 °Cd - in the leaf area (***A***), the internode diameter (***B***), and the internode length (***C***) according to organ relative rank, for rose primary axes grown under HH, LH, or LL in Exp. 1. The period between FBV stage and FBV + *ca*. 130 °Cd was characterized by sugar accumulation in HH, LH. The horizontal dashed line separates the upper expanding organs to the bottom fully-expanded ones. For each treatment, organs were pooled in 8 groups according to their relative position (see M&M) from plant base. For each group, data are the mean ± SEM of the differences between values at FBV + *ca*. 130 °Cd and the mean value at FBV (*n*= 4-5 plants). Different letters indicate significant differences between treatments (ANOVA followed by a Tukey test, *p* < 0.05).

### Transient light limitation induces sugar accumulation due to reduced organ expansion, while continuous light limitation reduces it through low photosynthesis

The combined estimations of photosynthesis and measurements of organ expansion (Fig. 5,6) support the hypothesis that the reduced sugar accumulation observed in LL after FBV results from limited photosynthesis and thus reduced C supply. Conversely, the sugar over-accumulation in LH appears to result from reduced organ expansion, leading to lower C demand while C supply remains similar to HH. This interpretation relies on two assumptions (i) in LL, the reduction in C demand by growing organs does not compensate for the reduction in C supply; (ii) in LH, the reduction in C demand is sufficient to increase the supply–demand ratio relative to HH. Testing these assumptions requires a quantitative comparison of C supply and demand dynamics.

We estimated the temporal dynamics of C demand of all primary-axis organs every 10 °Cd from FBV to 300 °Cd (end of the sugar accumulation phase; Fig. 3) as the C required for structural growth. Structural C growth was calculated as the product of organ expansion (leaf area increase or internode/peduncle volume expansion) and structural C content per unit dimension (area, volume), with structural C defined as total C minus soluble C compounds. Leaf expansion and internode/peduncle elongation were estimated using a previously developed rose model (Demotes-Mainard *et al*., 2013a) fitted to the data from Exp. 1 (Fig. S10-S11). Internode and peduncle diameters, leaf structural mass per unit area, and internode structural mass per unit volume were directly fitted on Exp. 1 measurements (Fig. S12-14).

From these estimates, total leaf expansion, internode elongation, and to a lesser extent internode thickening were found to be higher in HH than in LH or LL, consistent with Fig. 6, while internode thickening was higher in LH than in LL (Fig. 7A). Leaf structural mass per unit area remained nearly constant in LL and HH, but increased slightly in LH, suggesting a possible investment in structures during expansion following the return to high light (Fig. S13). Internode structural mass increased similarly across treatments and did not contribute to treatment differences in demand (Fig. S14). Structural growth of roots, the flower, and secondary axes was included in total demand but did not differ significantly between treatments (data not shown).

**Figure 7.**
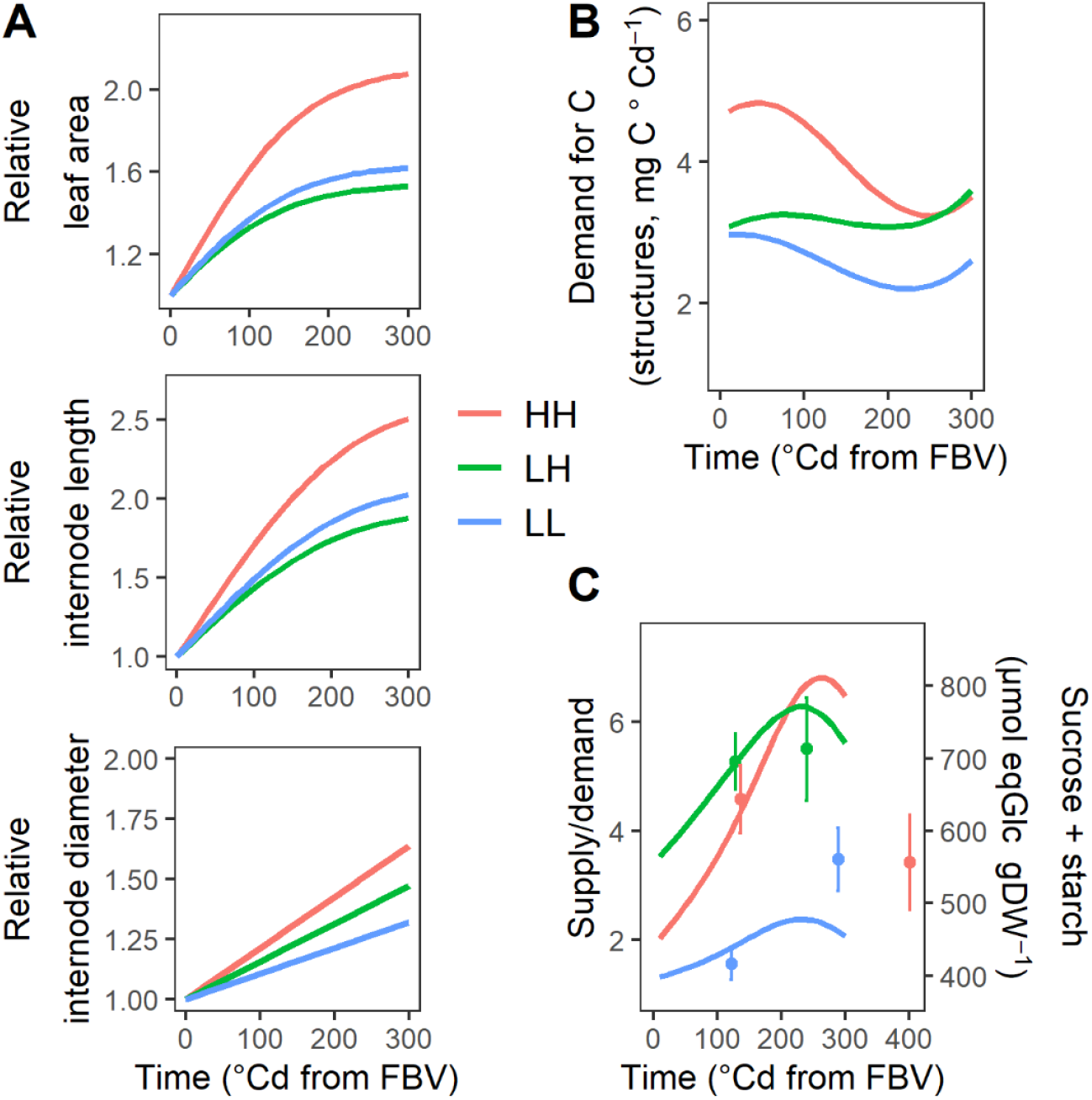
Continuous and temporary light limitation reduced C demand after FBV, and respectively repressed and stimulated C supply-demand balance just after FBV. (***A***) Estimated temporal dynamics - from FBV till 300 °Cd - of components for structural mass estimation of primary axes of Exp. 1 rose plants under HH, LH, or LL: leaf areas, internode/peduncle lengths and diameters. Data on individual organs were cumulated and expressed relatively to values at 0°Cd for lisibility. Estimations for individual organs are presented Fig. S10 to S14. (***B***) Derived estimated temporal dynamics of C demand. Derived supply-demand ratio and measured sucrose+starch internode contents (Z2+Z3). Sugar values were means ± SEM (*n*=4-5 plants; data similar to Fig. 3). Methods for estimations are detailed in material and methods. The simulated primary axes had 9, 8, 8 leaves for HH, LH, LL, respectively, according to the median total number of leaves in each light treatment.

C demand was then calculated as the increment in total structural C between consecutive 10 °Cd steps after FBV (Fig. 7B). Immediately after FBV, HH displayed the highest C demand, approximately 1.5 times that of LH and LL. C subsequently decreased in HH and LL as organ expansion slowed (Fig. 7A). In LH, C demand started at values similar to LL at FBV, reflecting reduced organ expansions under both treatments, and remained relatively stable over time, reaching HH levels only around 200 °Cd. This indicates that C demand was lower in LH than in HH during the period of sugar accumulation observed in LH.

We next calculated the supply–demand balance by dividing photosynthetic C supply (Fig. 5D) by the estimated C demand (Fig. 7C). From FBV till *ca*. 250 °Cd, both HH and LH showed increasing supply due to expanding photosynthetic area (Fig. 5A, C), resulting in an increasing supply-demand balance consistent with the accumulation of sucrose and starch (Fig. 3A). Compared to HH, LH displayed a higher balance until *ca*. 200 °Cd primarily due to reduced demand, whereas LL consistently displayed a lower balance as a result of reduced photosynthesis. These treatment-specific patterns matched the observed sucrose and starch concentrations (Fig. 7C, symbols).

Together, these results quantitively validate our initial hypothesis: under continuous light limitation, reduced sugar levels are explained by limited photosynthesis, whereas under transient light limitation, enhanced sugar storage relative to HH results from reduced organ expansion and the associated decline in C demand.

### C supply-demand balance contributes to establishing bud outgrowth differences between treatments

We hypothesized that light-induced differences in the C supply-demand balance, and the resulting differences in sugar availability, underlie the observed differences in bud outgrowth. To test this, in a fourth experiment (Exp. 4), C balance was manipulated by either reducing supply in LH or increasing supply in HH and LL.

In LH, C supply was reduced by either masking all fully expanded leaves of the primary axis after FBV or applying DCMU, a photosynthesis inhibitor. Leaf masking significantly reduced bud outgrowth frequencies in both the intermediate (Z2) and basal (Z3) zones (Fig. 8A). DCMU application also repressed bud outgrowth, although the effect was weaker and restricted to Z3 (Fig. 8B).

**Figure 8.**
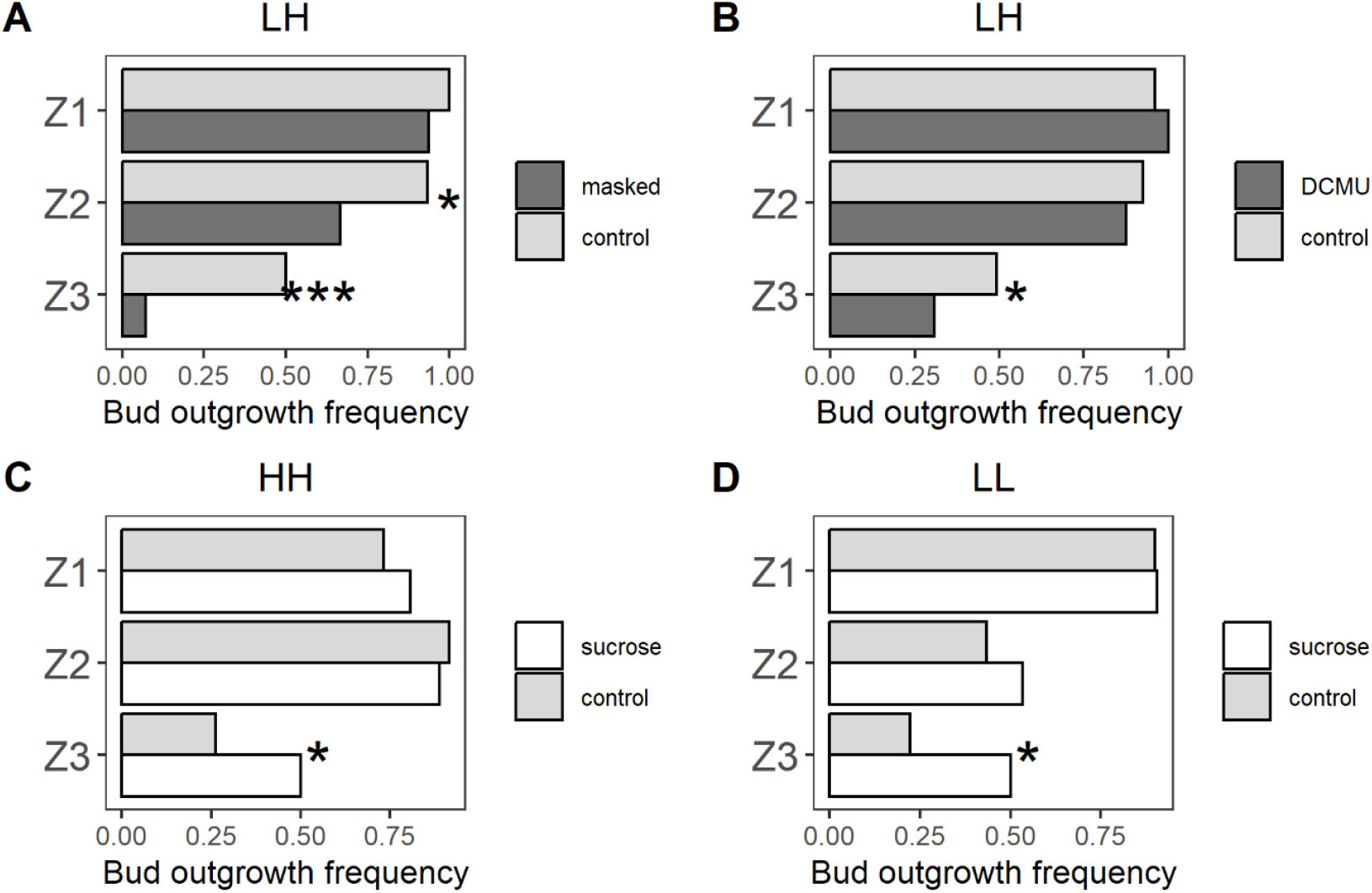
Changes of C supply induced disruptions in bud outgrowth response to light. (***A, B***) Bud outgrowth rate in the three zones along rose primary axes grown under LH with photosynthesis either not decreased (control) or decreased: (*A*) by covering fully opened leaves with either a black plastic (masked) or a transparent film (control) (*B*) or by applying 100 μM DCMU. (***C***,***D***) Bud outgrowth rate in the three zones along rose primary axes grown under HH (*C*) or LL (*D*) and supplied with either 50 mM mannitol (control) or 100 mM sucrose in zone Z3 of the primary axis. Bar represent the mean number of outgrowing buds over each zone 20 days after FBV (*n*=15, 20, 9, 15 for *A, B, C, D*, respectively). Asterisks indicate significant difference between bud outgrowth rates of controlled and treated plants (Fisher’s exact test; *: *p*<0.05; ***: *p* <0.001).

In HH and LL, C supply was increased by delivering exogenous sucrose through the petiole of the upper Z3 leaf after FBV. Mannitol was used as an osmotic control and, and an additional sucrose source was supplied above the application site to maintain sucrose delivery to upper sink organs. In both HH and LL, petiole-fed sucrose significantly stimulated bud outgrowth in Z3 compared with mannitol (Fig. 8C, D).

Together, these results demonstrate that treatment-dependent differences in bud outgrowth are causally linked to light-induced variations in the C supply–demand balance and the associated sugar availability.

## Discussion

Bud outgrowth is strongly influenced by light intensity, yet the mechanisms underlying this effect remain unclear (Schneider *et al*., 2019). In particular, the long-standing hypothesis-implemented in some ecophysiological models-that light-mediated control of bud outgrowth involves changes in sugar availability has never been directly tested. Although the promotive role of sugar availability in bud vicinity and related-signalling pathways is well-established (Barbier *et al*., 2015a; Mason *et al*., 2014), sugar availability involvement in light responses relies on correlative evidence and has been questioned in favour of cytokinins (CK) (Corot *et al*., 2017; Roman *et al*., 2016a). Furthermore, the mechanisms underlying the effects of transient low-light episodes remain unclear. Using a quantitative and dynamic analysis of source-sink relationships in single-axis rose plants, combined with source-sink perturbations, our study reveals that continuous and transient light limitation have contrasting effects on bud outgrowth-respectively inhibitory and stimulatory-through differential impacts on carbon (C) source and sink functions. The resulting changes in sugar availability within the plant axis contribute causally to bud outgrowth responses to light regimes.

### Continuous and transient light limitation have contrasting effects on C availability, mainly through differential impacts on source and sink activities

Reduced light intensity is classically associated with a decline in plant C status, reflected by lower sugar contents in organs or leaf mass per unit area (Bertin and Gary, 1998; Corot *et al*., 2017; Furet *et al*., 2014; Lafarge *et al*., 2010; Proietti *et al*., 2023; Savvides *et al*., 2014; Tardieu *et al*., 1999), consistent with the strong dependence of photosynthesis on irradiance (e.g., Hirose and Werger, 1987; Tardieu *et al*., 1999). In agreement with this framework, continuous low light reduced sucrose and starch concentrations in the median and basal parts of the rose primary axis (Fig. 3). In contrast, the effects of transient light limitation are much less documented. Here, a short period of low light followed by a return to high light induced a strong increase in sucrose and starch immediately after the light shift, with starch levels reaching values up to six times higher than under continuous high light (Exp. 2, Fig. 3). Similar responses have been reported in rice leaves and sheaths following transient shading (Lafarge et al., 2010).

Because light intensity affects not only photosynthesis but also induces morphological and physiological adaptations (Chenu *et al*., 2005; Grindlay, 1997; Jiang *et al*., 2021), we conducted an integrated analysis to disentangle the respective contributions of C sources and sinks to plant C status. C supply was estimated from individual leaf area and surfacic photosynthesis, driven by local light and photosynthetic capacity, while C demand was approximated from structural growth. As expected, continuous low light strongly reduced surfacic photosynthesis (Fig. 5, 7B, 7C). This was accompanied by increased area of mature basal leaves, typical of shade-avoidance (Demotes-Mainard *et al*., 2016) (Fig. 5A), and reduced expansion of upper organs (Fig. 6). Although these morphological responses slightly increased C supply and reduced demand, they were insufficient to compensate for the limitation of surfacic photosynthesis, resulting in markedly reduced whole-axis photosynthesis compared with continuous high light.

Our analysis also provides an explanation for the counterintuitive sugar accumulation observed after transient light limitation. This accumulation resulted from a non-reversible inhibition of upper organ expansion induced under low light, which persisted after the return to high light (Fig. 5A, 6) and reduced C demand. This finding is consistent with observations showing that transient shading during early leaf development irreversibly limits final leaf size by reducing the initial cell pool (Granier and Tardieu, 1999a). Similar effects likely extend to monocots, where low light during the enclosed phase of leaf development delays entry into rapid elongation and reduces final leaf size (Muller *et al*., 2001). In our study, this irreversible effect extended to internodes, whose length and diameter were reduced by transient low light (Fig. 6B,C). Lower fully-expanded leaves were unaffected, likely because their early development occurred before the light treatments, as they were preformed in the bud of the original cutting (Girault *et al*., 2010).

In contrast to C demand, C source capacity after transfer to high light was only slightly altered relative to continuous high light (Fig. 5D). Reduced upper leaf area decreased self-shading and enhanced light interception by lower leaves, while limiting photosynthesis of upper leaves (Fig. 5B,C). These compensatory effects resulted in similar whole-axis photosynthesis between transient and continuous high light conditions. Transient low light also induced non-reversible reductions in leaf photosynthetic capacity (Fig. 4C,D), consistent with previous reports (Oguchi *et al*., 2005; Sims and Pearcy, 1992) but in our conditions, these physiological adaptations has a limited impact on whole-axis photosynthesis (Fig. 5C).

### C availability as a determinant of light-mediated bud outgrowth regulation

Evidence linking C availability to environmentally driven branching responses has mainly emerged from cereal studies, based on correlations between tiller number and indicators of source–sink balance under varying light and temperature conditions (Alam *et al*., 2014; Alam *et al*., 2017; Kim *et al*., 2010). In rose, biochemical approaches have provided more direct support for such a relationship (Corot *et al*., 2017; Furet *et al*., 2014). Here, we show that transient low light strongly stimulated bud outgrowth, whereas continuous low light inhibited it relative to continuous high light (Fig. 2) in parallel with differences in sucrose and starch levels (Fig. 3). Our analysis further revealed that, under continuous low light, bud outgrowth was delayed and occurred only after sufficient sugar accumulation in stems, suggesting that a minimal C availability is required to trigger bud activation. This is consistent with *in vitro* studies showing that a threshold sugar supply is required to overcome auxin-mediated inhibition of bud outgrowth (Bertheloot *et al*., 2020). In addition, under transient low light, the interval between the outgrowth of successive buds along the axis was shortened (Fig. 2B), coinciding with strong sugar accumulation. This acceleration explains the over-stimulation of bud outgrowth and branching at axis scale (Fig. 2). A similar reduction in lag time before bud outgrowth has been reported for rose buds *in vitro* under increased sugar availability (Barbier *et al*., 2015a).

Recent studies have questioned the involvement of sugars in light-mediated bud regulation, as exogenous sugar supply failed to restored branching in rose plants transferred to darkness or low light (Corot *et al*., 2017; Roman *et al*., 2016a). Our results challenge this conclusion. Under transient low light condition, reducing C source through leaf masking or photosynthetic inhibition inhibited bud outgrowth, while under continuous low light, exogenous sugar supply promoted it (Fig. 8). Differences with previous studies likely arise from experimental conditions and sugar application methods. In Roman et al. (2016), plants were decapitated, defoliated, and placed in darkness, which may have activated specific signalling pathways (Yoshida *et al*., 2011). In Corot et al. (2017), very high sucrose level was supplied through the petiole (300–800 mM) (Bertheloot *et al*., 2020; Mason *et al*., 2014) which may have induced inhibitory effects, and vascular feeding (50-200 mM) may have preferentially supplied apical sinks rather than buds. In contrast, our combined medium petiole and vascular feeding strategy (100 mM sucrose) (Fig. S1) enabled moderate sugar supply to the bud while satisfying apical demand, thereby promoting bud outgrowth.

Both Corot et al. (2017) and Roman et al. (2016) identified CK as mediators of light effects on bud outgrowth, as CK levels decreased under low light and exogenous CK restored branching. Similar conclusions were drawn for wheat tillering (Lorenzo *et al*., 2015). I*n vitro* studies further show that light or CK act synergistically with sugar availability to stimulate bud outgrowth : light or CK alone, without sugar, cannot induce bud outgrowth, whereas sucrose enhanced CK effect for buds under light, including in presence of auxin (Bertheloot *et al*., 2020; Girault *et al*., 2010; Jiang *et al*., 2025; Rabot *et al*., 2012). This synergistic effect involves a stimulation of CK signalling, bud metabolism and sink activity by sugar (Jiang *et al*., 2025). Thus, two interacting pathways may coexist in light-mediated bud regulation : a sugar- and a CK-dependent pathway. Such combined regulation has also been reported for shoot apical meristems (Li *et al*., 2017; Pfeiffer *et al*., 2016).

### Key model requirements for simulating bud outgrowth regulation at the whole-plant scale

Many whole-plant models simulate organ growth—and sometimes branch initiation—as a function of plant C status, often through a C supply–demand balance (e.g. Drouet and Pages, 2007; Evers *et al*., 2010; Guo *et al*., 2006; Lescourret *et al*., 2011; Luquet *et al*., 2006; Pallas *et al*., 2013). Our study highlights requirements for models aiming to simulate branching responses to light intensity, including transient low-light conditions.

Models must explicitly account for light history effects on organ development. Transient exposure to low light during early developmental stage strongly limits subsequent organ expansion, even after return to high light (Fig. 6,S8; Cookson *et al*., 2005; Granier and Tardieu, 1999a), leading to reduced C demand (Fig. 7A,B). When C supply is only weakly affected, as observed in our study, this creates a decoupling between C supply (high) and demand (reduced), resulting in sugar over-accumulation and enhanced bud outgrowth. In addition, C demand must be formalized as a function of organ expansion rather than as an increase in mass, which also includes changes in dry mass par unit organ dimension. The latter is linked to plant C status and so reflects the outcome of the source-sink balance rather than the intrinsic demand of growing organs (Hilty *et al*., 2021; Tardieu *et al*., 1999). These requirements may extend to the modelling of plant responses to water deficit, which similarly limits leaf expansion, can increase plant C status, and was reported to stimulate branching upon stress release (Blum *et al*., 1990; Demotes-Mainard *et al*., 2013b; Granier and Tardieu, 1999b; Lecoeur *et al*., 1995).

A major challenge, not addressed in the present study, is the formalization of bud outgrowth regulation by C availability. Current models rely on empirical relationships between C status and branching, implemented either at the whole-plant scale or at the level of individual buds with predefined windows of bud sensitivity (Evers *et al*., 2010; Hammer *et al*., 2023; Luquet *et al*., 2006). However, these formalisms have rarely been evaluated against detailed experimental datasets and do not capture how buds dynamically integrate C availability over time. Physiological evidence indicate that sugars act not only as substrates but also as signaling molecules, activating within the bud pathways involving trehalose-6-phosphate and hexokinase, sugar transporters, vacuolar invertase activity—a marker of sink strength—, and metabolic pathways (Barbier *et al*., 2021; Fichtner *et al*., 2017; Fichtner *et al*., 2020; Henry *et al*., 2011; Wang *et al*., 2021). Sugar signaling is also tightly interconnected with hormonal regulation: high sugar availability represses strigolactone signaling and promotes CK signaling, leading to repression of the dormancy-associated gene BRC1 (Bertheloot *et al*., 2020; Patil *et al*., 2022)(Cao *et al*., 2023; Jiang *et al*., 2025)(Mason *et al*., 2014; Wang *et al*., 2019). Integrating this mechanistic knowledge into a bud-scale model would represent a key step toward a physiologically grounded representation of bud regulation by C availability. While Bertheloot et al (2020) proposed a model describing the sugar–hormone interaction network controlling bud outgrowth, further developments are now required to dynamically represent bud outgrowth for integration within dynamic whole-plant models of branching plasticity.

## Conclusion

Using a quantitative and dynamic modelling approach that captures source–sink responses to light regimes and source–sink manipulations, we confirm that continuous low light intensity inhibits bud outgrowth through reduced photosynthesis. We identify, for the first time, the mechanism underlying the non-intuitive stimulatory effect of transient light limitation: a reduction in the growth—and thus C use —of apical sink organs appearing under low light (Fig. 9). This mechanism may extend to other environmental factors limiting sink growth, such as water deficit. Accurately simulating branching plasticity therefore requires models that accurately represent both photosynthesis and the regulation of C use by sink organs. This, in turn, calls for a better quantitative understanding of how bud outgrowth regulators are dynamically integrated at the bud level. In addition to C, CK contribute to light-mediated bud outgrowth and likely act as a complementary regulatory pathway. How C- and CK-mediated controls interact remains an open question. Beyond light intensity, light quality—modulated by canopy structure and controlled-environments—also affects branching and its interaction with light quantity represents an important avenue for future research.

**Figure 9.**
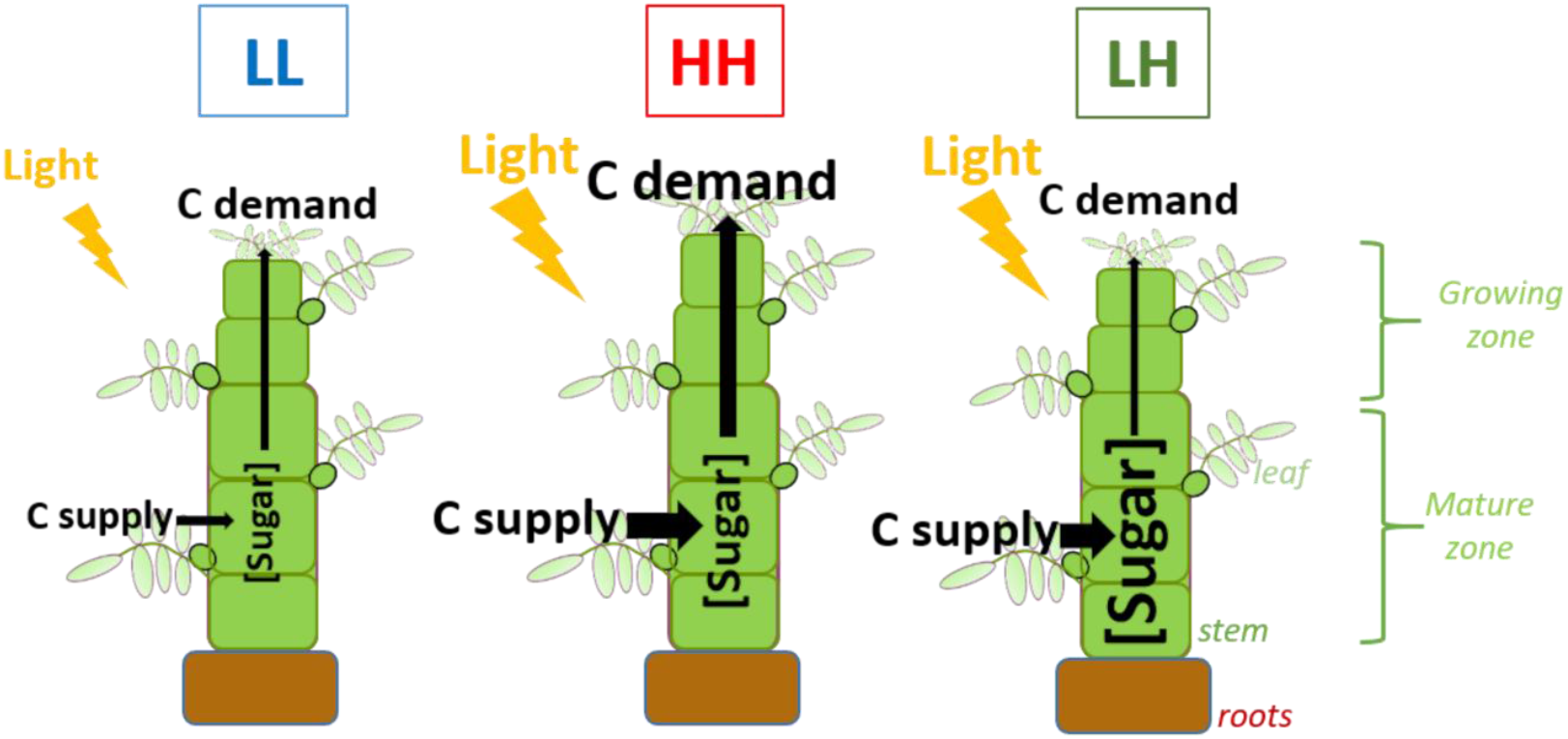
Schematic diagram of the effect of light treatment on the source–sink balance within the plant. Compared to plants submitted to continuous high light (HH), plants submitted to continuous low light (LL) have a lower sugar availability due to reduced incident light (Fig. 5B) inducing a lower photosynthesis and carbon (C) supply (Fig. 5D); the development of apical organs is also reduced, inducing a reduction in C demand (Fig. 7B), but this effect was much smaller than the effect on C supply, explaining the lower sugar availability. Conversely, plants submitted to transient low light (LH)-ending just before the start of bud outgrowth period-, display a higher sugar availability due to a non-reversible reduction of apical organ development (Fig. 6) and thus C demand (Fig. 7B), while C supply by photosynthesis was poorly affected (Fig. 5D).

## Supplementary data

Figure S1. Picture and scheme describing the method used for exogenous sugar supply.

Figure S2. Bud outgrowth along primary axes of rose plants grown under different light treatments in Exp. 2.

Figure S3. Bud outgrowth and dry mass dynamics under the different light treatments.

Figure S4. Increasing sugar availability for bud-bearing stem segments *in vitro* promotes sucrose and starch contents in the stem.

Figure S5. Leaf expansion periods for primary axes of plants grown in Exp. 3.

Figure S6. Final number of leaves in the different light treatments for Exp.1.

Figure S7. Measured PPFD at each leaf level after FBV for the different light treatments (Exp. 3).

Figure S8. Measured final organ dimensions for light treatments HH and LH (Exp. 3).

Figure S9. Variables used in equations 3-4 to estimate expansion kinetics of leaves and internodes for primary axes of rose plants in Exp. 1 under HH, LH, LL.

Figure S10. Simulated and measured area expansions of individual leaves for each light treatment.

Figure S11. Simulated and measured elongations of individual internode and peduncle for each light treatment.

Figure S12. Estimated and measured diameter increase of individual internodes and peduncle for each light treatment.

Figure S13. Estimated and measured structural mass per unit area for individual leaves for each light treatment.

Figure S14. Estimated and measured structural mass per unit volume for individual internodes and peduncle for each light treatment.

Figure S15. Estimated and measured chlorophyll index for individual leaves in each light treatment for Exp. 3.

## Acknowledgments

We thank Gérard Sintès for assistance with the growth chamber, Odile Douillet for help with plant growth, and Bénédicte Dubuc and the ImHorPhen team for rose production (UMR IRHS).

## Author contribution

AS: Conceptualization, Visualization, Investigation, Methodology, Data curation, Formal analysis, Software, Writing-original draft. FB: Methodology, Writing-review & editing. SDM: Writing-original draft, Writing-review & editing. LL: Investigation, Methodology, Validation. MDPG: Investigation, Methodology, Visualization. CC: Investigation, Methodology. YG: Methodology, Supervision, Writing-review & editing. CG: Supervision, Conceptualization, Methodology, Writing-review & editing. SS: Funding acquisition, Supervision, Conceptualization, Writing-original draft, Writing-review & editing. JB: Funding acquisition, Project administration, Supervision, Conceptualization, Methodology, Formal analysis, Software, Data curation, Visualization, Writing-original draft, Writing-review & editing.

