## Supplemental figures for "Branching Varies with Light Limitation Scenarios in relation with Changes in Carbon Source-Sink Dynamics"

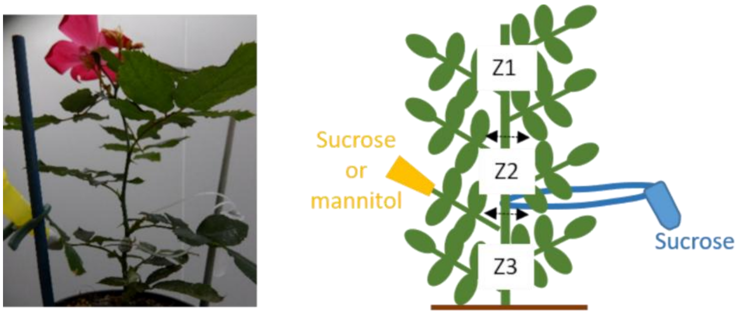


**Figure S1. Picture and scheme describing the method used for exogenous sugar supply.** Either sucrose (100 mM) or mannitol (50 mM) were supplied through the petiole of the topmost leaf of Z3 (in yellow). For both sugar treatments, 100 mM sucrose was also vascularly supplied (in blue) to satisfy sugar demand of the growing apical organs of the plant. The horizontal double arrows indicate the separation between the three pre-defined zone of the primary axis: Z1, Z2, Z3 (see material and methods for calculation details).

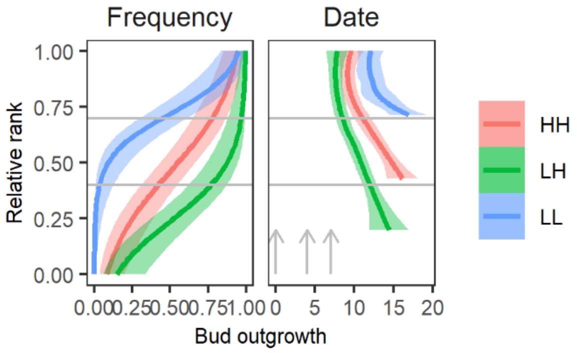


**Figure S2. Bud outgrowth along primary axes of rose plants grown under different light treatments in Exp. 2.** Outgrowth frequency and date (expressed as degree days after FBV) for buds at different ranks along the primary axis of plants grown under HH, LH, and LL in Exp. 2. Rank is expressed relative to the total number of leaves and assigned to a zone (Z1, Z2, Z3) as described in material and methods. Lines represent logistic (Frequency) and loess (Date) smooth lines ± 95% confidence intervals (*n*=15 plants for LH and 20 plants for HH, LL). For outgrowth date, only ranks with an outgrowth frequency above 0.3 are represented. Grey arrows represent sampling dates for sugar analysis (Fig. 3B).

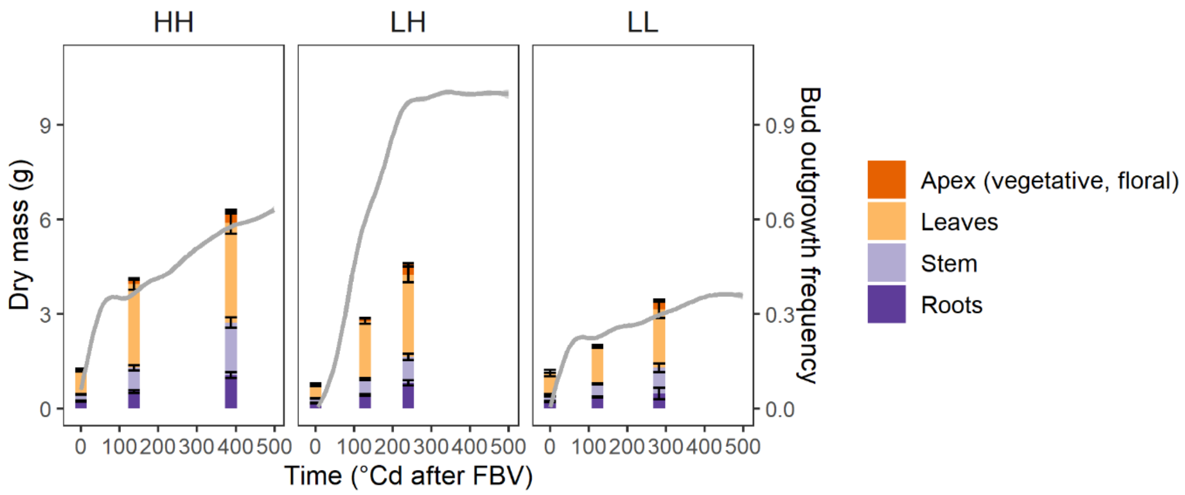


**Figure S3. Bud outgrowth and dry mass dynamics under the different light treatments.** Dry mass of the primary axis and roots, its distribution between organ types (stacked bars), and cumulative bud outgrowth frequency over the whole primary axis (grey lines) over time for plants grown under HH, LH, and LL in Exp. 1. Individual colored bars are means ± SEM (*n*=4-5 plants). Organs types distinguished for dry mass are the roots, the stem (internodes, nodes, peduncle), the leaves, and the apex (all apical organs that could not be individualized because too small). Grey lines represent loess smooth lines ± 95% confidence intervals.

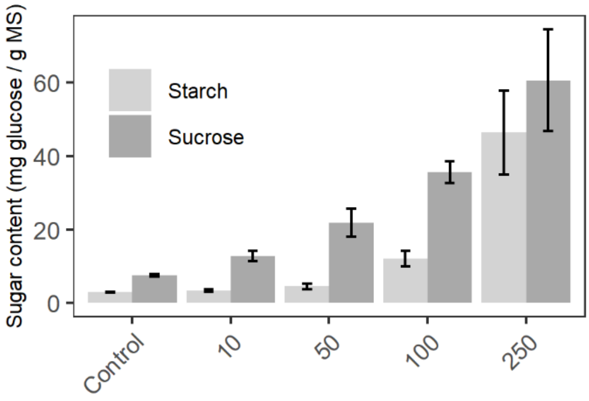


**Figure S4. Increasing sugar availability for bud-bearing stem segments *in vitro* promotes sucrose and starch contents in the stem.** Starch (light grey) and sucrose (dark grey) concentrations in the stem 48h after stem segments excision from primary axis of rose plants at FBV and placed in a culture medium with an osmotic control (mannitol), 10, 50, 100, or 250 mM sucrose. Data are means ± SEM (*n*=3 plants).

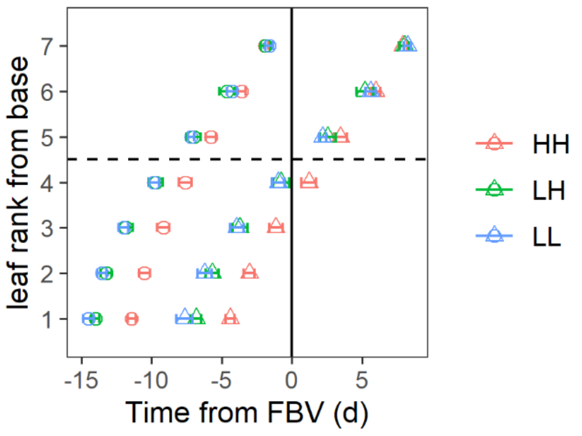


**Figure S5. Leaf expansion periods for primary axes of plants grown in Exp. 3.** Dates at which individual leaves appeared (circles) and reached 90% their final size (triangles) for the primary axis of rose plants grown under HH, LH, and LL. The vertical full line represents the date of FBV, the horizontal dashed line separates the bottom group of leaves that had ended their expansion at FBV and the upper one that expanded after FBV. Data are means ± SEM (*n*=15-17 plants).

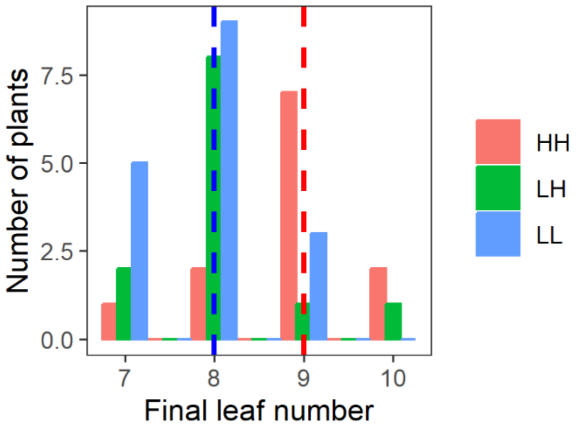


**Figure S6. Final number of leaves in the different light treatments for Exp.1.** Number of plants with 7, 8, 9, or 10 leaves formed on the primary axis for treatments HH (red), LH (green), and LL (blue) for the group of harvested plants in Exp. 1. The vertical red and blue dashed lines correspond to the median number of leaves for HH and LL, respectively (significantly different, Student's test: $p < 0.05$). The green line corresponding to LH is masked by the blue line.

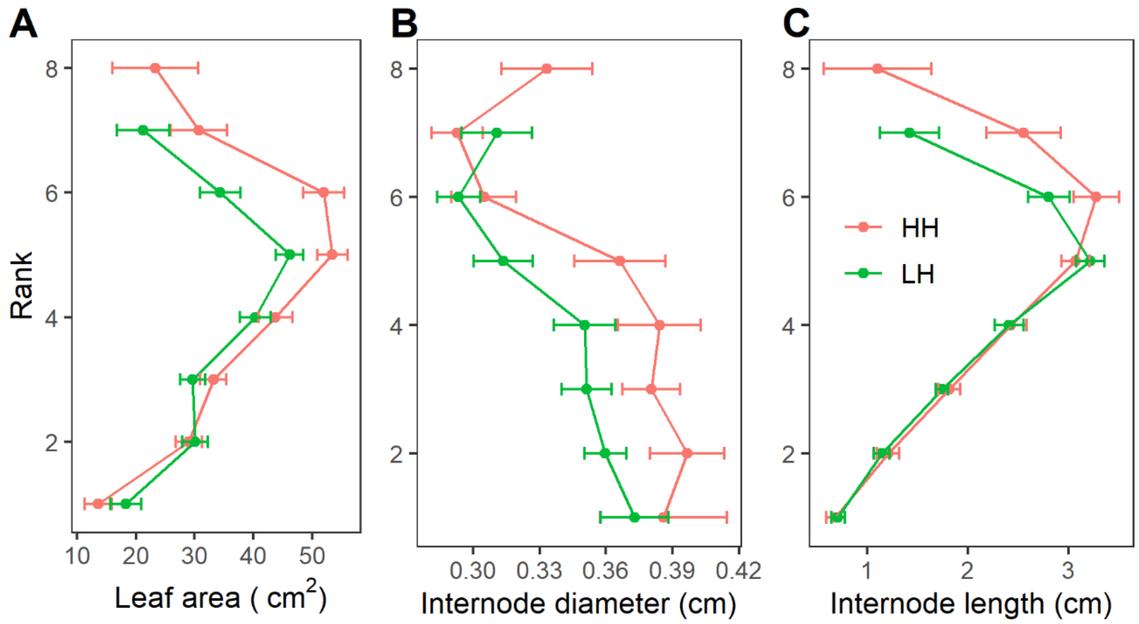


**Figure S7. Measured PPFD at each leaf level after FBV for the different light treatments (Exp. 3).** PPFD was measured at 3, 10, and 14 days after FBV at leaf rank levels 1, 3, 4 (counting from base) of primary axes of rose plants grown under HH, LH, or LL. Data represent mean ± SEM (*n*=10 plants). Asterisks indicate significant differences between LH and HH (Student's test: $p < 0.05$).

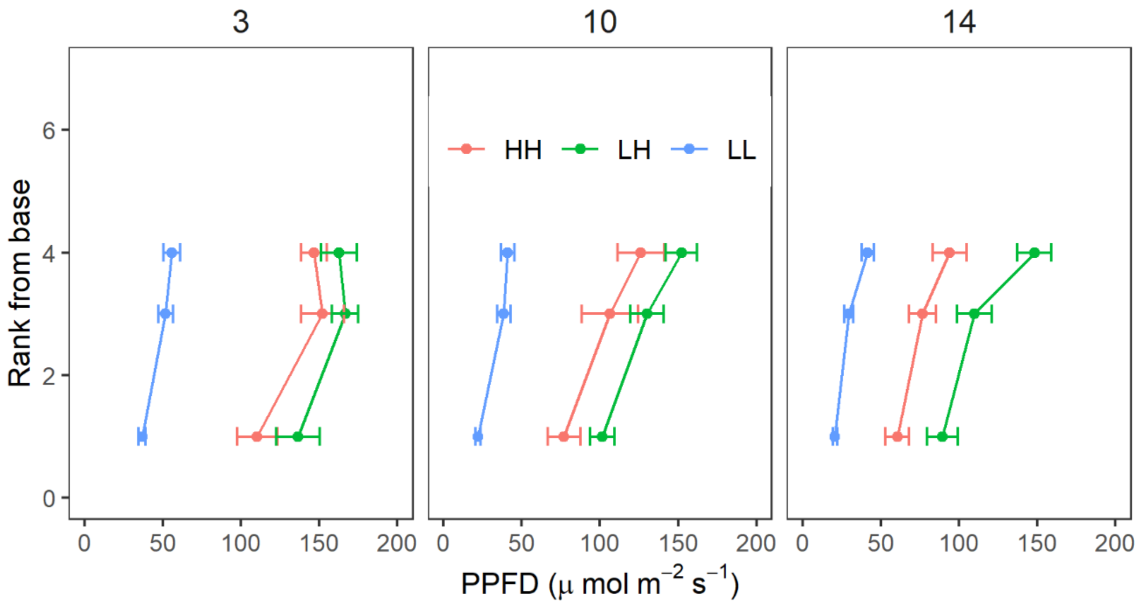


**Figure S8. Measured final organ dimensions for light treatments HH and LH (Exp. 3).** Final areas of leaves (*A*), diameters of internodes (*B*), and lengths of internodes (*C*) as a function of their rank from the base of primary axes of rose plants grown under HH or LH. Dimensions were measured at OF (opened flower stage). Data represent means ± SEM (*n*=14-15 plants). Asterisks indicate significant differences between means (Student's test: $p < 0.05$).

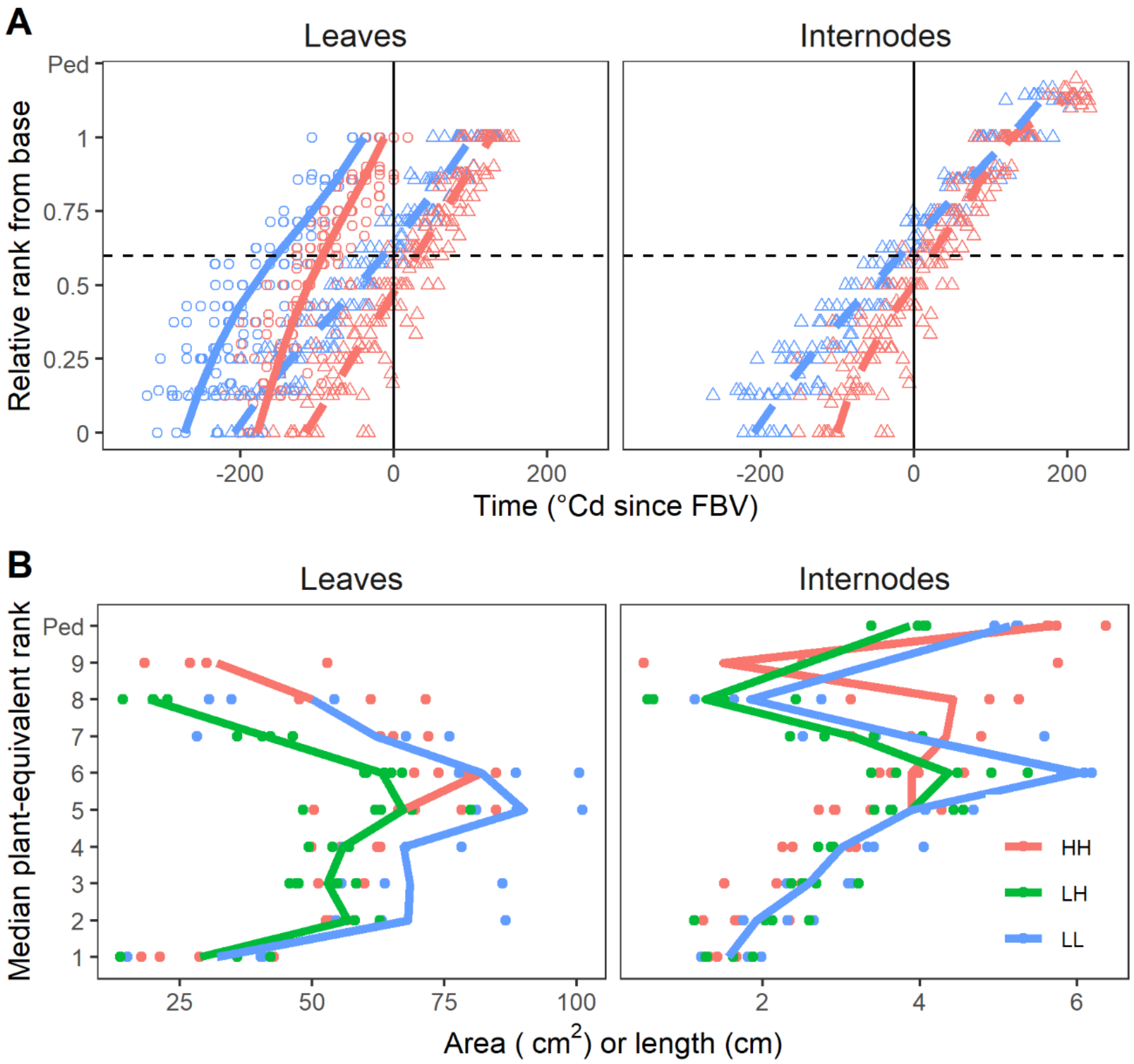


**Figure S9. Variables used in equations 3-4 to estimate expansion kinetics of leaves and internodes for primary axes of rose plants in Exp. 1 under HH, LH, LL.** (***A***) Dates of leaf appearance (full lines) and leaf and internode/peduncle maximum relative expansion rate ( $t^*_{i,lf}$ $t^*_{i,in}$ dashed lines) as a function of organ relative position. Circles represent observations of leaf appearance dates on individual plants (*n*= 21-22, non-destructive plant group), triangles represent the corresponded estimated dates of maximum expansion rates for leaves and internodes/peduncle (see M&M for details), lines represent the estimations (cubic smoothing spline) for a median plant in each light treatment. The vertical line corresponds to FBV, the horizontal dashed lines separates organs that have finished their expansion at FBV to those which extends after FBV. Data in LH were taken identical to those in LL. ***(B)*** Maximum leaf area ( $A^{max}_{i,lf}$ ) and internode or peduncle length ( $L^{max}_{i,in}$ ) as a function of organ rank from base. Symbols represent observations for individual plants (n=3-4 plants), lines the mean at each rank with treatments pooled when there was no significant difference (anova). Ranks are the median plant-equivalent ones calculated from relative ranks (see M&M for details). Maximum dimensions were taken on the last harvesting date for each light treatment (destructive plant group). *Ped*= Peduncle. Axis of median plants was made of 8 leafy phytomers for LL and LH, and 9 for HH.

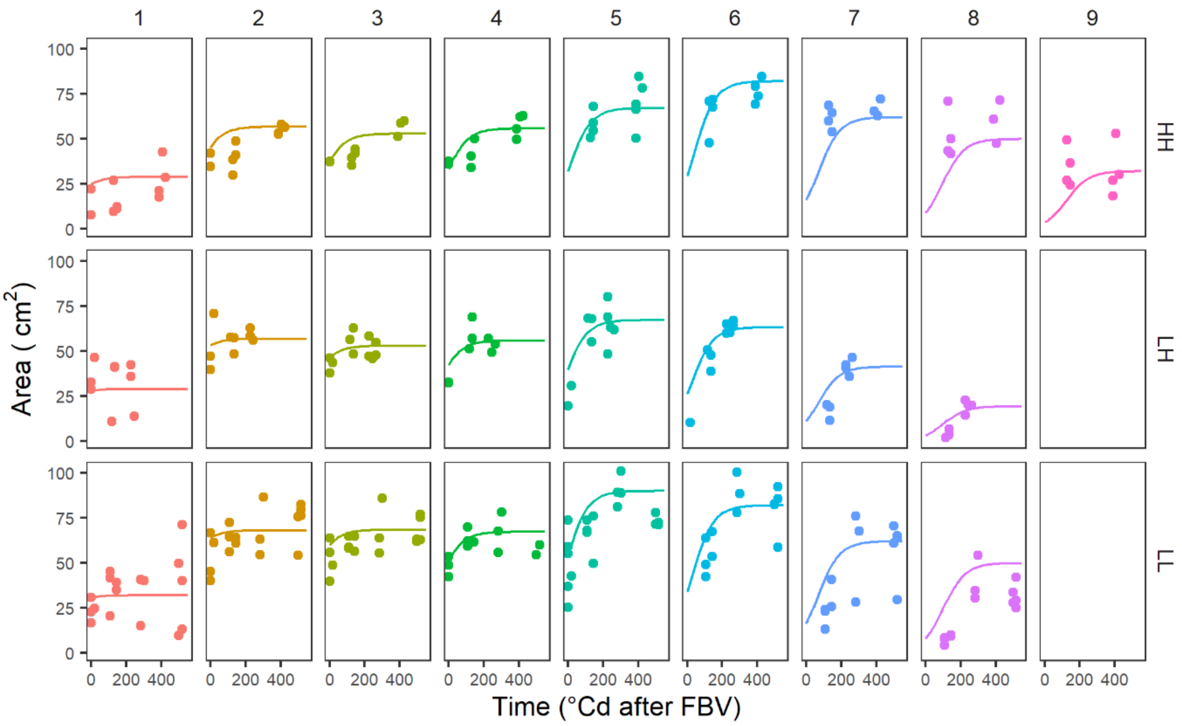


**Figure S10. Simulated and measured area expansions of individual leaves for each light treatment.** Simulations (lines) *vs.* measured (symbols) leaf area through thermal time from FBV for each rank (each graph column) of a primary axis of median plant in HH, LH, or LL. Simulated areas were obtained from equation (3) in which the times of maximum relative expansion rates and individual leaf final areas were estimated from measurements as described Fig. S9. Measured leaf areas correspond to individual plants of Exp. 1 (*n*=4-5 plants), and leaf ranks were recalculated and expressed as median plant-equivalent rank (see M&M for details) with median plants having 9 leaves in total on the primary axis in HH, and 8 leaves in LH and LL (Fig. S6).

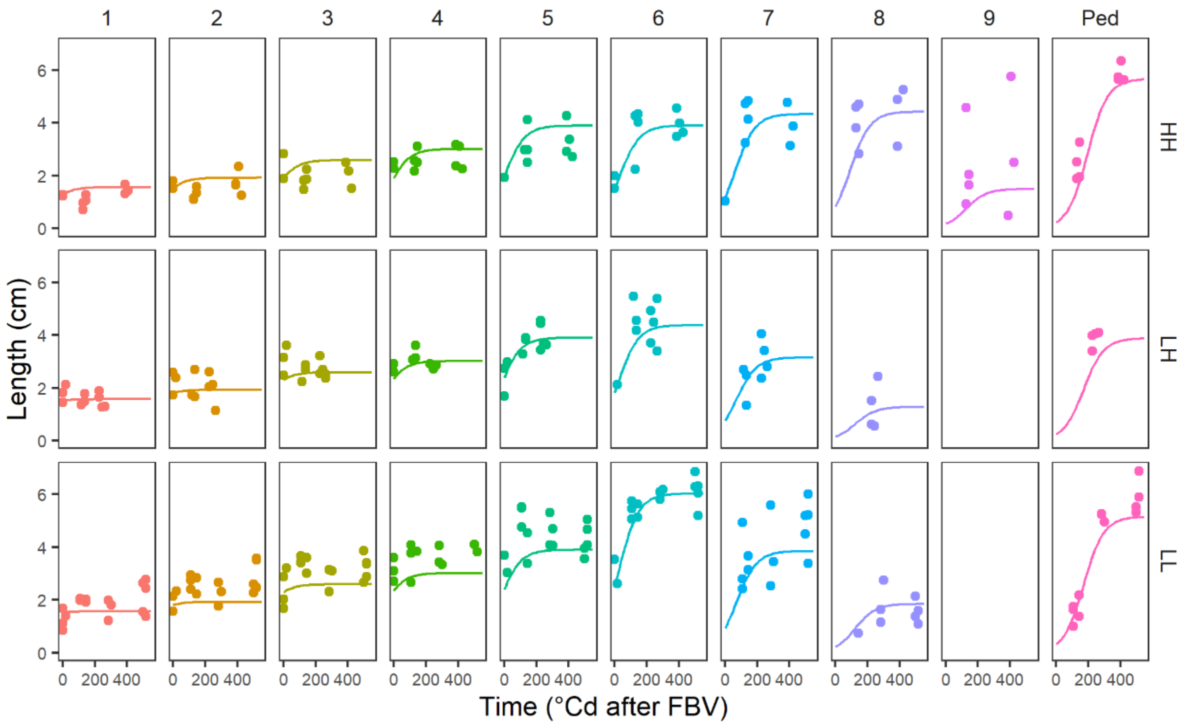


**Figure S11. Simulated and measured elongations of individual internode and peduncle for each light treatment.** Simulated (lines) *vs.* measured (symbols) internode/ peduncle length through thermal time from FBV for each rank (each graph column) of a primary axis of a median plant in HH, LH, or LL. Simulated lengths were obtained from equation (4) in which times of maximum relative expansion rates and individual internode and peduncle final lengths were estimated from measurements as described in Fig. S9. Measured lengths correspond to individual plants of Exp. 1 (*n*=4-5 plants), and leaf ranks were recalculated and expressed as median plant-equivalent rank (see M&M for details) with median plants having 9 leaves in total on the primary axis in HH, and 8 leaves in LH and LL (Fig. S6). *Ped*=peduncle.

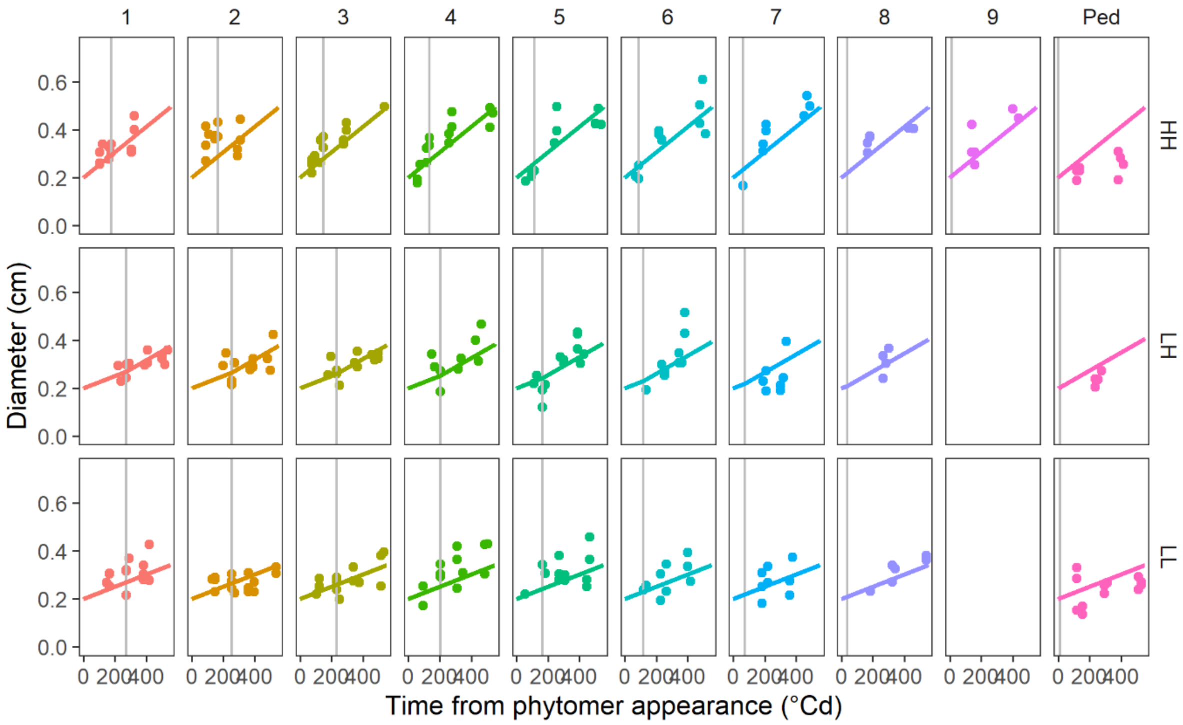


**Figure S12. Estimated and measured diameter increase of individual internodes and peduncle for each light treatment.** Estimated (lines) *vs.* measured (symbols) internode/peduncle diameters through thermal time from phytomer appearance date for each rank (each graph column) of a primary axis of a median plant in HH, LH, or LL. Vertical grey lines represent FBV date. Diameters were estimated from Exp. 1 data (see M&M for details). Measured diameters correspond to individual plants of Exp. 1 (*n*=4-5 plants), and leaf ranks were recalculated and expressed as median plant-equivalent rank (see M&M for details) with median plants having 9 leaves in total on the primary axis in HH, and 8 leaves in LH and LL (Fig. S6). *Ped*=peduncle.

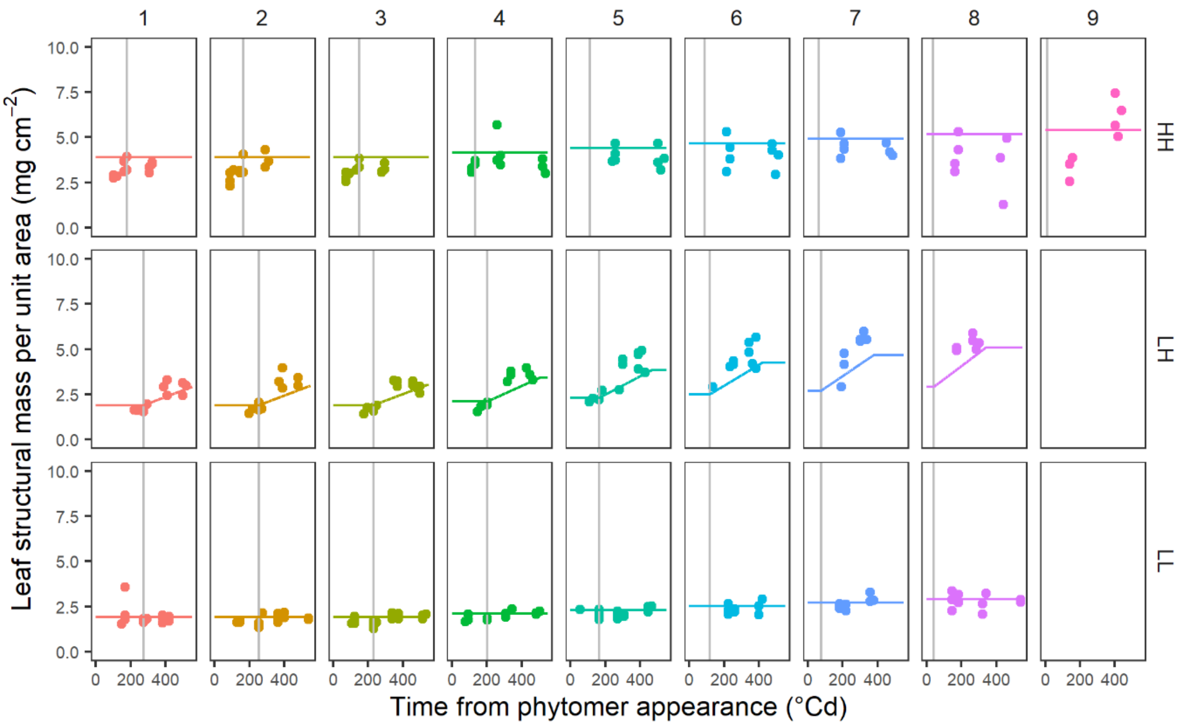


**Figure S13. Estimated and measured structural mass per unit area for individual leaves for each light treatment.** Estimated (lines) *vs.* measured (symbols) leaf structural mass per unit area through thermal time from leaf appearance date for each rank (each graph column) of a primary axis of a median plant in HH, LH, or LL. Vertical grey lines represent FBV date. Structural masses per unit area were estimated from Exp. 1 data (see M&M for details). Measured masses correspond to individual plants of Exp. 1 (*n*=4-5 plants), and leaf ranks were recalculated and expressed as median plant-equivalent rank (see M&M for details) with median plants having 9 leaves in total on the primary axis in HH, and 8 leaves in LH and LL (Fig. S6). *Ped*=peduncle.

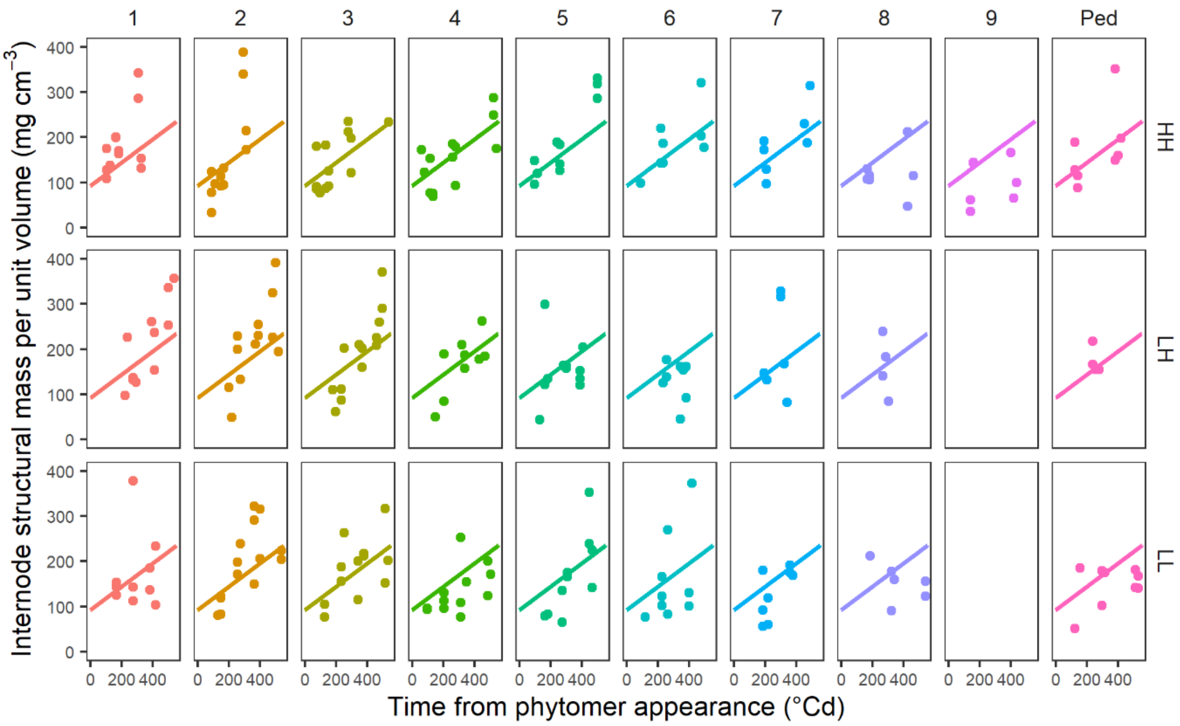


**Figure S14. Estimated and measured structural mass per unit volume for individual internodes and peduncle for each light treatment.** Estimated (lines) *vs.* measured (symbols) internode/peduncle structural mass per unit volume through thermal time from phytomer appearance date for each rank (each graph column) of a primary axis of a median plant in HH, LH, or LL. Vertical grey lines represent FBV date. Structural masses per unit area were estimated from Exp. 1 data (see M&M for details). Measured masses correspond to individual plants of Exp. 1 (*n*=4-5 plants), and leaf ranks were recalculated and expressed as median plant-equivalent rank (see M&M for details) with median plants having 9 leaves in total on the primary axis in HH, and 8 leaves in LH and LL (Fig. S6). *Ped*=peduncle.

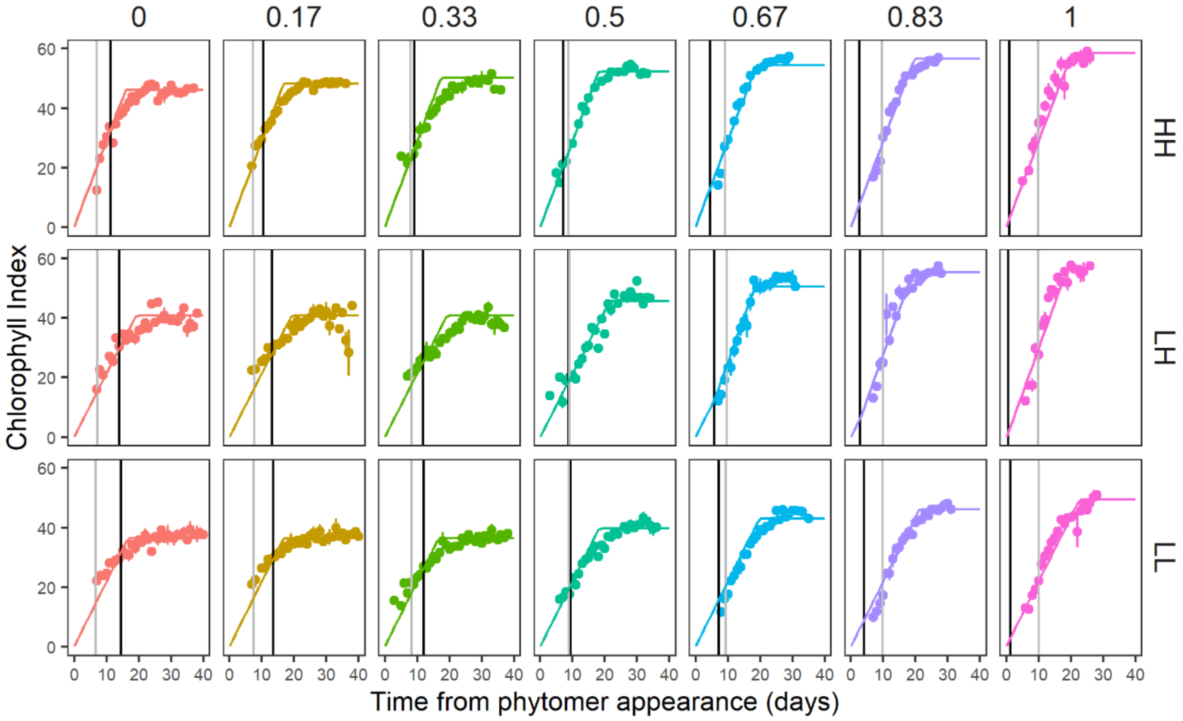


**Figure S15. Estimated and measured chlorophyll index for individual leaves in each light treatment for Exp. 3.** Estimated (lines) *vs.* measured (symbols) leaf chlorophyll index through time from leaf appearance date for each rank (each graph column) of a primary axis of a median plant in HH, LH, or LL. Vertical grey lines represent FBV date. Leaf chlorophyll indices were estimated from Exp. 1 data (see M&M for details). Measured indices correspond to mean ± SEM (*n*=18 plants), and leaf ranks were expressed as relative ranks from primary axis base (see M&M for details). Median plants have 7 leaves that had appeared in total on the primary axis.
